# Preimplantation factor (PIF) is an endogenous inhibitor of potassium channel K_V_1.3 regulating neutrophil function during pregnancy

**DOI:** 10.64898/2026.03.20.713251

**Authors:** Roland Immler, Wiebke Nadolni, Johanna M Franz, Annika Bertsch, Sebastian Baasch, Vasilios A Morikis, Aleksandra Kurova, Marco Borso, Ignasi Forné, Ericka C M Itang, Johannes B Müller-Reif, Monika Pruenster, Lou M Wackerbarth, Matteo Napoli, Ina Rohwedder, Anna Yevtushenko, Michael Rauer, Martin Kolben, Markus Moser, Eytan Barnea, Melanie Boerries, Thomas Vogl, Scott I Simon, Claudia Klein, Philipp Henneke, Axel Imhof, Susanna Zierler, Markus Sperandio

## Abstract

Pregnancy is a unique period regarding immune cell regulation. Within the placenta, maternal immune cells play a central role in immune surveillance and tissue remodeling. However, regulatory mechanisms of systemic immunity during pregnancy are less clear. Here, we show that neutrophil function is altered in pregnant mice (E13.5), indicated by increased slow rolling velocity and reduced adhesion. Mechanistically, PreImplantation factor (PIF), a 15 amino acid peptide which is produced by human and murine trophoblast cells of the placenta, is continuously secreted into the maternal circulation and plays a key role in modulating neutrophil function via blocking the voltage-gated potassium channel K_V_1.3. This resulted in impaired intracellular Ca^2+^ signaling and subsequently disturbance of neutrophil post-arrest modifications and a higher susceptibility to physiological shear forces *in vivo* and *in vitro*. Furthermore, PIF-mediated K_V_1.3 blockade impaired E-selectin-mediated release of S100A8/A9 and phagocytosis. Taken together, we have identified PIF as an important modulator of neutrophil function during pregnancy suggesting a critical role in regulating innate immune responses throughout gestation.

## Introduction

Pregnancy is a unique and delicate period that requires tight immune cell regulation. On one hand, the maternal immune system has to protect the mother and the fetus from internal/external threats. At the same time, maternal immune cells require tight control in order to avoid immune-mediated damage to the growing fetus and termination of pregnancy. Nevertheless, the concept that pregnancy reflects a constant period of systemic and local immune suppression has recently been challenged by demonstrating that alternating pro- and anti-inflammatory periods are a prerequisite for successful parturition ^1^. Implantation, placentation and later parturition are characterized by a pro-inflammatory signature, accompanied by immune cell infiltration into the decidua and placenta ^2^. An immune-regulatory environment identifies the intermediate period of fetal growth and development ^3^. These alternating pro- and anti-inflammatory stages are largely mediated locally by the endometrium, the decidua and the placenta together with immune cells present in the respective tissues ^1^. In early pregnancy, endometrial stromal cells and immune cells secrete for example interleukin-6 (IL-6), CXCL8 (IL-8; human), CXCL1 (mouse) or TNF which further attracts immune cells and promotes trophoblast invasion ^4^. In addition, invading immune cells release pro-angiogenic factors such as vascular endothelial growth factor (VEGF) or placental growth factor (PGF), allowing vascularization and tissue remodeling of the decidua ^5^. Both, trophoblast cells and immune cells then start to produce a variety of primarily anti-inflammatory mediators and growth factors interfering with cytotoxicity of immune cells and shaping local immune tolerance ^6, 7, 8^.

A number of pregnancy-specific factors have been linked to maternal immune modulation, including the hormones progesterone or estradiol ^9, 10^. Recently, a 15 amino acid peptide named PreImplantation factor (PIF) has been proposed to have immune modulatory functions during pregnancy, too ^11^. PIF is expressed and secreted by trophoblast cells ^12, 13^ and can be detected in maternal circulation at varying levels throughout pregnancy ^14, 15^. Sufficient serum concentration of PIF is required for embryo development and successful birth ^15, 16^. Properties of PIF regulating immune cell function have been broadly addressed, including in several models for autoimmune diseases outside pregnancy ^17, 18^. However, the molecular mode of action of PIF on immune cells has not been elucidated.

Neutrophils account for 70% of the total population of circulating leukocytes in human peripheral blood. They are the first cells recruited to sites of inflammation and are fully equipped to fight off against invading pathogens or clear cell debris ^19^. Their recruitment to sites of inflammation follows a well-defined cascade, initiated by tethering and rolling along the inflamed vascular endothelium ^20^. Chemokines and alarmins, immobilized on the endothelium or released from neutrophils trigger the activation of β_2_ integrins and concomitant cell arrest ^21^. Finally, neutrophils crawl along and transmigrate through the endothelium to enter the inflamed tissue ^22^. Moreover, neutrophils have been implicated to play an important role in many autoimmune diseases ^21, 23^.

Here, we set out to investigate the role of trophoblast-derived peptide PIF on the function of maternal neutrophils during acute inflammation. We show that PIF inhibits the voltage-gated potassium channel K_V_1.3, thereby affecting neutrophil post-arrest modifications, S100A8/A9 release as well as phagocytic activity. Thus, we identify PIF as a highly potent endogenous anti-inflammatory agent and suggest a potential role of PIF as a therapeutic molecule beyond its function during pregnancy.

## Results

### Neutrophil recruitment is altered during pregnancy

Increased susceptibility to infections during pregnancy suggests altered immune cell function, especially altered innate immunity ^24^. To study neutrophil recruitment during pregnancy *in vivo*, we applied intravital microscopy (IVM) of TNF stimulated mesentery postcapillary venules in pregnant (E13.5) *Catchup^IVM-red^* mice, a neutrophil-specific reporter mouse line ^25^ (Fig. 1A). Analysis of neutrophil recruitment 3h post-stimulation revealed no differences in the number of rolling neutrophils along postcapillary venules in pregnant mice compared to non-pregnant, aged-matched controls (Fig. 1B). In contrast, neutrophil rolling velocity was significantly increased (Fig. 1C, D) and numbers of adherent neutrophils were significantly reduced in pregnant animals (Fig. 1E), revealing dampened neutrophil recruitment during pregnancy. Of note, hemodynamic parameters (vessel diameter, blood flow velocity, white blood cell (WBC) and neutrophil counts) did not differ among the groups (Supplemental Table S1).

**Figure 1.**
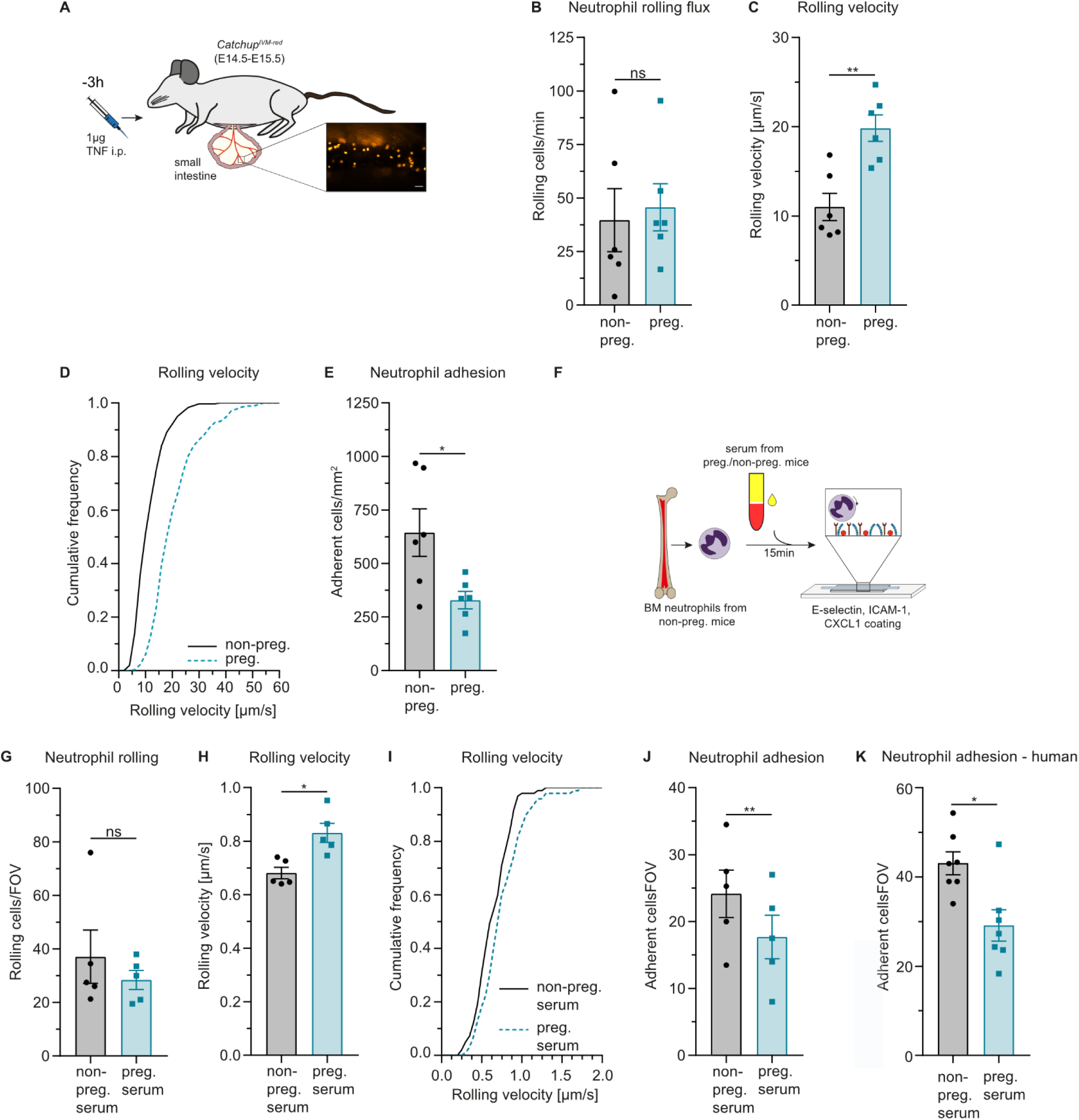
Neutrophil recruitment is altered during pregnancy. (**A**) Experimental design of IVM of the mouse mesentery. (**B**) Rolling flux (number of rolling cells), (**C**) mean rolling velocities, (**D**) cumulative distribution of rolling velocities and (**E**) neutrophil adhesion was assessed in pregnant and non-pregnant *Catchup^IVM-red^*mice treated with 1µg TNF (i.p.) 3h prior to investigation (n=6 mice per group, unpaired Student’s t-test; cumulative frequency: n=382 (non-preg) and n=316 (preg) cells). (**F**) Experimental design of flow chambers with serum incubation. (**G**) Number of rolling cells (**H**) mean rolling velocities, (**I**) cumulative distribution of rolling velocities and (**J**) number of adherent neutrophils incubated with serum from pregnant and non-pregnant mice perfused through E-selectin/ICAM-1/CXCL1-coated flow chambers (n=5 mice per group, paired Student’s t-test; cumulative frequency: n=97 (non-preg serum) and n=98 (preg serum) cells). (**K**) Number of adherent neutrophils incubated with human serum from pregnant and non-pregnant individuals perfused through E-selectin/ICAM-1/CXCL8-coated flow chambers (n=7 independent experiments, paired Student’s t-test). Data is presented as mean±SEM and as cumulative frequency.

Analysis of surface molecules on peripheral blood neutrophils, which impact recruitment revealed significantly higher expression levels of CD11b (Mac-1, α_M_), CD62L (L-selectin) and CD162 (PSGL-1) in pregnant *WT* mice (Supplemental Figure S1) which is in line with elevated CD11b and CD62L levels on peripheral neutrophils from pregnant women ^26^. Since this pro-inflammatory surface marker signature of neutrophils from pregnant individuals did not match with reduced neutrophil recruitment observed, we hypothesized that factors in the serum might alter neutrophil function during pregnancy. To investigate this in more detail, we perfused E-selectin, ICAM-1 and CXCL1-coated flow chambers with bone marrow neutrophils isolated from non-pregnant (male and female) *WT* mice which had been incubated for 15min with serum from pregnant or non-pregnant mice (Fig. 1F). In line with our *in vivo* findings, pre-incubation of the cells with serum from pregnant dams did not alter the number of rolling neutrophils (Fig. 1G), but increased rolling velocities (Fig. 1H,I) and reduced cell adhesion to the flow chamber substrate (Fig. 1J). This indicates that factors in the serum of pregnant mice alter the ability of neutrophil to roll and adhere along/at inflamed endothelium within 15min of incubation. To compare the dampening effect of pregnant serum in humans, we conducted flow chamber experiments with neutrophils from male donors preincubated with serum from pregnant women (week 20-25 of gestation). In line with the observations in mouse cells, presence of serum from pregnant women decreased neutrophil adhesion under shear flow compared to those incubated with non-pregnant serum (Fig. 1K). Together, these findings suggest pregnancy-associated serum factors that dampen neutrophil recruitment *in vivo* and *in vitro*.

### PreImplantation factor (PIF) impairs neutrophil recruitment *in vivo* and *in vitro*

The recently described pregnancy-specific small peptide PIF has been proposed to have immune modulatory functions during pregnancy ^11^. Therefore, we analyzed how PIF affects neutrophil recruitment *in vivo* using IVM of TNF-stimulated post-capillary venules of the mouse cremaster. To do so, we injected 1µg of synthetic (s)PIF into the scrotum (i.s.) of male WT mice 1h prior to the onset of inflammation (Fig. 2A). As controls, either a scrambled version of PIF (scrPIF) or the carrier substance alone (Ctrl) was applied. Pretreatment with sPIF significantly increased rolling velocities along inflamed postcapillary venules (Fig 2B,C) and reduced stable neutrophil adhesion to inflamed endothelium compared to controls (Fig. 2D). Importantly, hemodynamic parameters did not differ among the groups (Supplemental Table 2S) and expression levels of recruitment relevant surface molecules (CD11a, CD11b, CD18, L-selectin, PSGL-1, CXCR2 and CD44) were not altered upon i.s. sPIF injection (Supplemental Fig. S2A-H). In addition, we analyzed neutrophil transmigration into inflamed cremaster tissue (Fig. 2E). sPIF significantly reduced the number of extravasated neutrophils compared to controls (Fig. 2F), demonstrating impaired neutrophil recruitment in the presence of PIF *in vivo*. In order to study whether PIF alters neutrophil recruitment by directly interfering with neutrophils, we performed flow chamber assays perfusing E-selectin, ICAM-1 and CXCL1-coated glass capillaries with whole blood from *WT* mice incubated with sPIF or vehicle for 15min (Fig. 2G). We found that sPIF increased rolling velocities and reduced neutrophil adhesion to the coated surface compared to control (Fig. 2H-J), indicating that the observed impairment of neutrophil recruitment *in vivo* was mediated via a direct effect of PIF on neutrophils.

**Figure 2.**
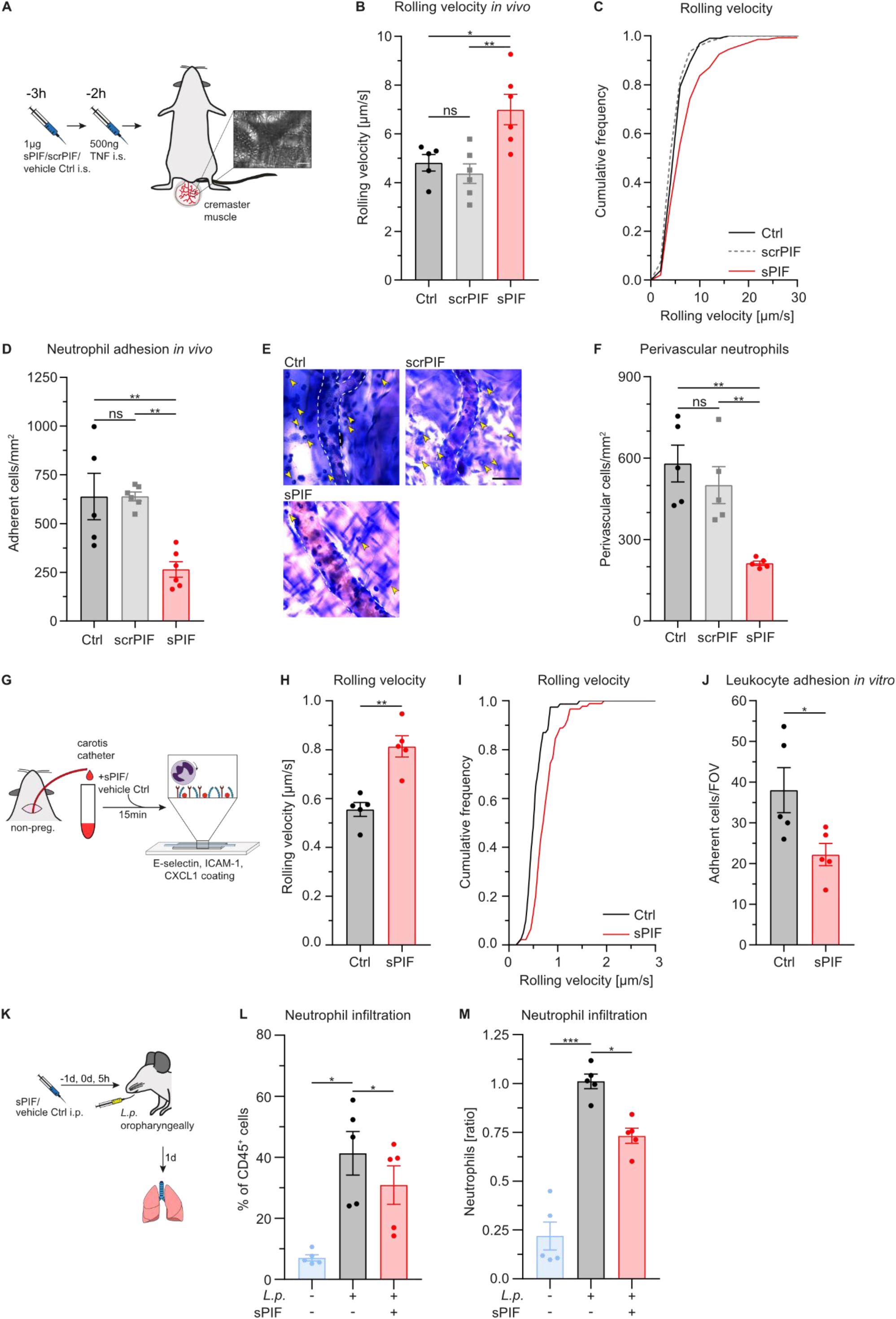
PIF impairs neutrophil recruitment *in vivo* and *in vitro*. (**A**) Experimental design of IVM of the TNF-stimulated mouse cremaster muscle. (**B**) Mean rolling velocities, (**C**) cumulative distribution of rolling velocities and (**D**) neutrophil adhesion was assessed in postcapillary venules of TNF stimulated (i.s.) cremaster muscles in *WT* mice pre-treated (i.s.) with 1µg sPIF, scrPIF or vehicle control (n=6 mice per group, one-way ANOVA, Tukey’s multiple comparison; cumulative frequency: n=101 (Ctrl), n=175 (scrPIF) and n=147 (sPIF) cells). (**E**) Stimulated cremaster muscles were stained with Giemsa (representative micrographs, scale bar: 30µm, yellow arrows: perivascular neutrophils) and (**F**) neutrophil extravasation was analyzed (n=5 mice per group, one-way ANOVA, Tukey’s multiple comparison). (**G**) Experimental design of flow chamber assays with murine whole blood. (**H**) Mean rolling velocities, (**I**) cumulative distribution of rolling velocities and (**J**) number of adherent neutrophils incubated with 300nM sPIF or vehicle control and perfused through E-selectin/ICAM-1/CXCL1-coated flow chambers (n=5 mice per group, unpaired Student’s t-test; cumulative frequency: n=78 (Ctrl) and n=90 (sPIF) cells). (**K**) Experimental design of respiratory tract infection with *L.p.*-dsRed. (**L**) Relative quantification and (**M**) Ratio of relative neutrophil numbers after L.p. infection with and without administration of 1µg sPIF i.p. at d-1, d0 and 5h post infection (pi) (n=5 mice per group; 1-way RM ANOVA; Dunnet’s multiple comparison). Data is presented as mean±SEM, as representative micrographs and as cumulative frequency.

To investigate the role of PIF on neutrophil behavior in a clinically relevant setting, we established a respiratory infection model using oropharyngeal instillation of *Legionella pneumophilia* (*L.p.*). To determine the optimal timing for sPIF administration, we initially monitored neutrophil recruitment in the first 8 hours post infection (pi). Neutrophil infiltration was detectable as early as 2h post infection (pi), followed by a marked increase at 4h pi that continued to rise through 8h (Supplemental Fig. S2I). As expected, monocyte recruitment was not significantly altered throughout this time course. Based on the observed kinetics, we injected 1µg sPIF i.p. at d-1, d0 and 5h pi to *L.p.*-infected *WT* mice and analyzed neutrophil recruitment to the lungs 1d pi (Fig. 2K). Administration of sPIF reduced neutrophil infiltration to the lungs by ∼25% compared to vehicle control injection (Fig. 2L, M), highlighting the capacity of PIF to modulate neutrophil recruitment *in vivo*. In addition, *L.p.* instillation resulted in weight loss 1d pi, which was less pronounced in PIF treated animals (Supplemental Fig. S2J).

Taken together, these results show that PIF reduces neutrophil recruitment *in vivo* and *in vitro* by directly interfering with neutrophil adhesion.

### Endogenous PIF alters neutrophil adhesion *in vitro*

Based on the neutrophil-modulating properties of sPIF *in vivo* and *in vitro*, we speculated that altered neutrophil recruitment in presence of serum from pregnant mice (Fig. 1H-K) is mediated via endogenous PIF. To address this hypothesis, we generated an antibody against PIF (clone 852/F2) and first tested its blocking properties *in vitro*. Accordingly, we perfused E-selectin, ICAM-1 and CXCL1 coated flow chambers with bone marrow neutrophils from non-pregnant *WT* mice pretreated with sPIF and the PIF antibody (Fig. 3A). Administration of PIF antibodies together with sPIF resulted in significantly reduced neutrophil rolling velocities and increased neutrophil adhesion compared to isotype control or sPIF alone (Fig. 3B-D), demonstrating that the PIF antibody neutralizes the effect of PIF on neutrophil recruitment *in vitro*. Importantly, addition of PIF antibodies without exogenous PIF did not alter neutrophil recruitment *in vitro* (Supplemental Fig. S3A-D), ensuring that the antibody did not directly activate neutrophils. Next, we incubated bone marrow neutrophils from non-pregnant *WT* mice with serum from pregnant mice in the presence of PIF antibodies and analyzed neutrophil recruitment in E-selectin, ICAM-1 and CXCL1 coated flow chambers (Fig. 3E). Addition of PIF antibodies reduced neutrophil rolling velocities (Fig. 3F,G) and increased neutrophil adhesion compared to isotype controls (Fig. 3H), suggesting that endogenous PIF in serum from pregnant mice impairs neutrophil recruitment *in vitro*. In line with this, addition of sPIF to whole blood from pregnant mice (Supplemental Fig. S3E) did not further increase neutrophil rolling velocities (Supplemental Fig. S3F,G), nor reduce adhesion in flow chamber assays (Supplemental Fig. S3H). These results demonstrate that endogenous PIF alters neutrophil recruitment *in vivo* and *in vitro*.

**Figure 3.**
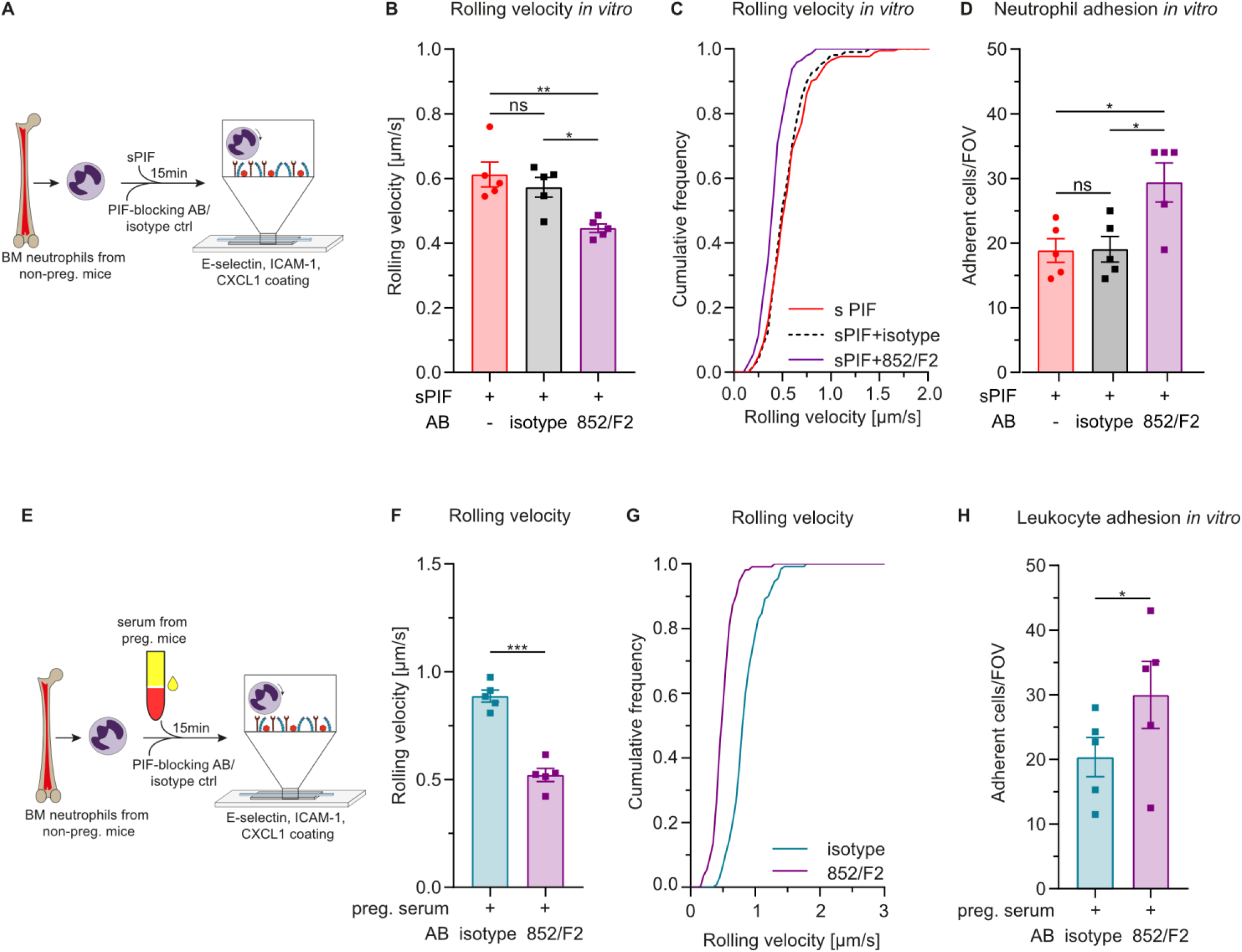
Endogenous PIF impairs neutrophil recruitment *in vitro*. (**A**) Experimental design assessing the PIF-blocking properties of antibody, clone 852/F2 in E-selectin, ICAM-1 and CXCL1 coated flow chambers. (**B**) Mean rolling velocities, (**C**) cumulative distribution of rolling velocities and (**D**) number of adherent neutrophils incubated with sPIF (300nM) and PIF-blocking antibody or isotype control (both 60µg ml^−1^) for 15min (n=5 independent experiments; one-way ANOVA, Tukey’s multiple comparison; cumulative frequency: n=171 (w/o antibody), n=107 (isotype) and n=145 (852/F2) cells). (**E**) Experimental design of flow chamber assays with PIF-blocking antibody. (**F**) Mean rolling velocities, (**G**) cumulative distribution of rolling velocities and (**H**) number of adherent neutrophils incubated with serum from pregnant mice and PIF-blocking antibody or isotype control (both 60µg ml^−1^; n=5 independent experiments, paired Student’s t-test; cumulative frequency: n=130 (isotype) and n=140 (852/F2) cells). Data is presented as mean±SEM, and as cumulative frequency.

### PIF is an endogenous inhibitor of the voltage-gated potassium channel K_V_1.3

Previously, mass spectrometry of decidual cells revealed a broad spectrum of putative interaction partners of PIF, including the voltage-gated potassium channel K_V_1.3 ^27^. Recently, we demonstrated that K_V_1.3 regulates multiple neutrophil functions *in vivo* and *in vitro* ^28, 29^. We therefore speculated that PIF interacts with K_V_1.3 thereby modulating neutrophil function. To address this, we first overexpressed human K_V_1.3 in HEK-293 cells and performed patch-clamp experiments. As expected, characteristic K_v_1.3 currents developed in response to voltage stimulation, reaching their maximum at +40mV (Fig. 4A, black lines). Notably, application of 300nM of sPIF almost completely abolished current activation in response to voltage stimulation (red lines). Quantification showed a characteristic increase in current-voltage relationships in response to increasing membrane voltages in control cells that was significantly inhibited in the presence of sPIF (Fig. 4B). Varying concentrations of sPIF applied to active K_v_1.3 currents revealed a dose-dependent inhibition of the channel with an IC_50_ of ∼10.2nM (Fig. 4C,D). To verify the inhibitory properties of PIF on K_v_1.3 channels on primary neutrophils, we applied the same consecutive step-protocol to isolated human neutrophils to induce K_v_1.3 currents. Cells exhibited typical voltage-dependent currents that were decreased by application of sPIF (Fig. 4E,F). In addition, application of sPIF on already active K_v_1.3 currents triggered via a single +40mV step again showed a strong inhibition of current amplitudes (Fig. 4G). This inhibition was even stronger than that observed in the presence of the specific small molecule inhibitor of K_V_1.3 PAP-1 (10nM) ^30^ (Fig. 4H).

**Figure 4.**
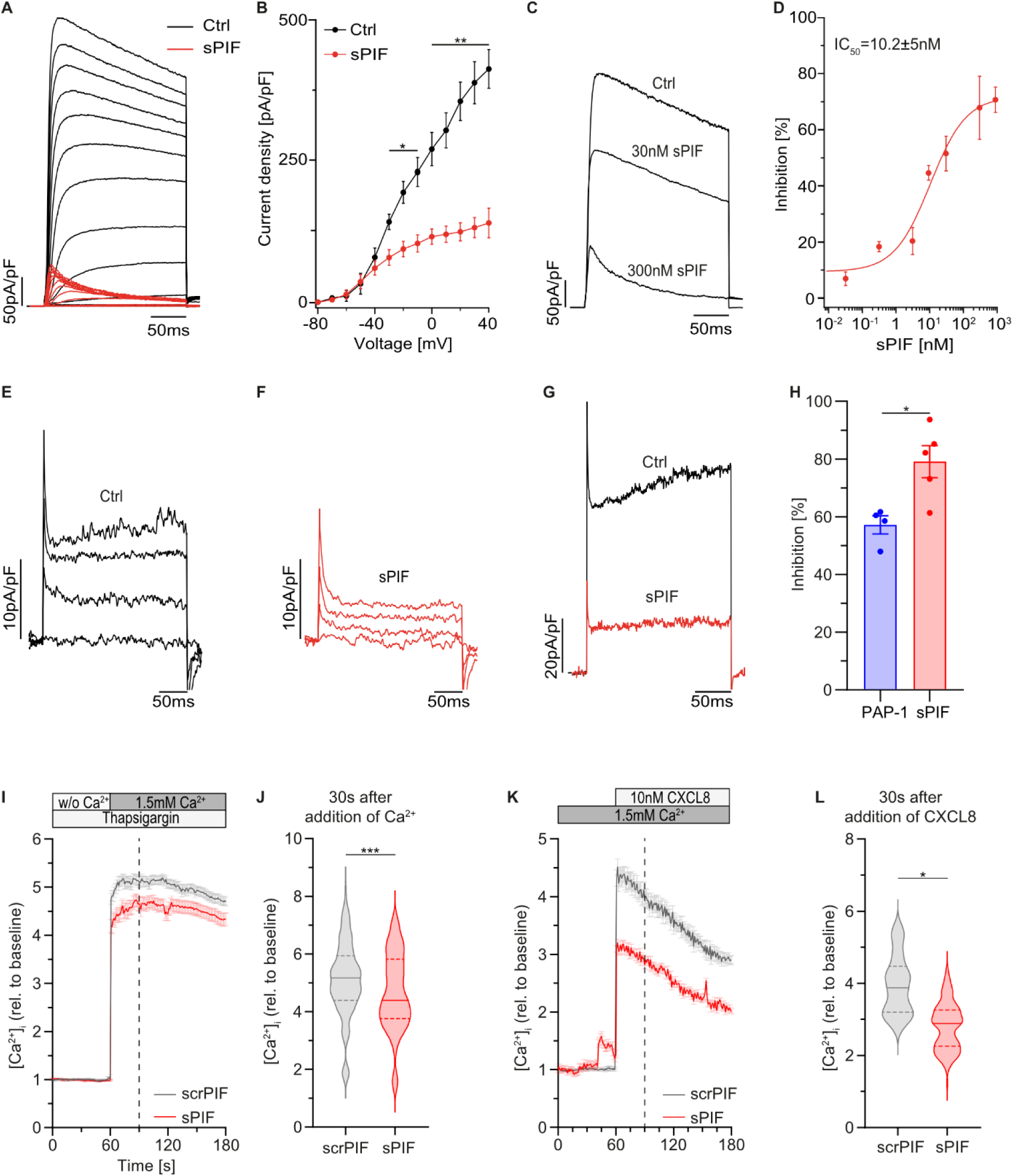
PIF is an endogenous inhibitor of the voltage-gated potassium channel K_V_1.3. (**A**) K_v_1.3 currents were measured using patch clamp in HEK-293 cells transiently overexpressing hK_v_1.3 (hK_v_1.3-HEK-293) by application of 13 consecutive 10mV steps from −80mV to +40mV over 200ms in absence (Ctrl) and presence of sPIF (300nM) (n=36 (Ctrl) and (n=4 (sPIF) cells, representative traces normalized to cell size as current density (pA/pF). (**B**) Current-voltage relationship of sPIF and control treated hK_v_1.3-HEK-293 cells (n=4 cells, repeated unpaired Student’s t-tests). (**C**) K_v_1.3 currents were extracted at +40mV in absence and after application of different concentrations of sPIF (representative traces). (**D**) Dose-dependent inhibition of K_v_1.3 currents (average peak current inhibition in %) in hK_v_1.3-HEK-293 cells by increasing concentrations of sPIF application was determined (n=4-8 cells, IC_50_=10.2±5nM. Hill=0.8±0.4). (**E,F**) K_v_1.3 currents in primary human neutrophils were induced by application of consecutive voltage-steps, (**E**) before (Ctrl) and (**F**) after addition of 300nM sPIF (representative traces no. 1, 4, 8, 13 from n=5 cells). (**G**) K_v_1.3 currents in primary human neutrophils were induced by the application of a single voltage-step to +40mV, before (Ctrl) and after application of 300nM sPIF (representative traces of n=5 cells). (**H**) Current inhibition [%] of K_v_1.3 currents in primary human neutrophils after application of the K_V_1.3-specific blocker PAP-1 (10nM) and sPIF (300nM) (n=4-5 cells, one-way ANOVA, Tukey’s multiple comparison). (**I**) CRAC channel dependent Ca^2+^ influx in Rhod-2 AM-loaded human neutrophils pretreated with Thapsigargin and 300nM sPIF or scrPIF was determined by addition of 1.5mM Ca^2+^. (**J**) [Ca^2+^]_i_ was quantified 30s after addition of Ca^2+^ (n=126 (scrPIF) and n=88 (sPIF) cells from 3 independent experiments, unpaired Student’s t-test). (**K**) Total Ca^2+^ flux was investigated after CXCL8 stimulation under static conditions. (**L**) Total [Ca^2+^]_i_ relative to baseline was quantified 30s after addition of CXCL8 (n=37 (scrPIF) and n=114 (sPIF) cells from 3 independent experiments, unpaired Student’s t-test). Data is represented as representative trances and as mean±SEM.

In lymphocytes and neutrophils, K^+^ efflux via K_V_1.3 helps to maintain the membrane potential thereby allowing sustained Ca^2+^ influx via store-operated calcium entry (SOCE) ^28, 31^. To test whether PIF-mediated inhibition of K_V_1.3 affects Ca^2+^ signaling, we monitored changes in intracellular Ca^2+^ concentrations ([Ca^2+^]_i_) in human neutrophils loaded with the Ca^2+^ indicator Rhod-2 AM. First, cells were pre-treated with the Sarco/Endoplasmic Reticulum Calcium ATPase (SERCA) inhibitor Thapsigargin in Ca^2+^ free conditions to induce endoplasmic reticulum Ca^2+^ store depletion and ensuing opening of CRAC channels. Addition of Ca^2+^ to the medium induced an increase in [Ca^2+^]_i_ in control cells which was significantly decreased in neutrophils incubated with sPIF (Fig. 4I,J). Next, we quantified GPCR-dependent Ca^2+^ signaling by stimulating the cells with CXCL8. Inhibition of K_V_1.3 using sPIF resulted in a significant reduction of [Ca^2+^]_i_ compared to controls (Fig. 4K,L), demonstrating that PIF treatment altered Ca^2+^ signaling in human neutrophils. Together, these results show that PIF is an endogenous inhibitor of K_V_1.3 thereby interfering with Ca^2+^ signaling in neutrophils.

### PIF-mediated inhibition of K_V_1.3 impairs post-arrest modifications, S100A8/A9 release and phagocytosis in neutrophils

Recently, we have shown that high [Ca^2+^]_i_ at LFA-1 cluster sites is a prerequisite for sufficient post-arrest modification steps in neutrophils (integrin outside-in signaling) ^32^, a process that requires K_V_1.3 activity ^28^. Inhibition or genetic deletion of K_V_1.3 resulted in impaired cytoskeletal rearrangement under flow conditions and a higher susceptibility of adherent neutrophils to physiological shear forces^28^. Therefore, we investigated whether K_V_1.3 inhibition via PIF influences cytoskeletal rearrangement, adhesion strengthening, and firm adhesion. To do so, we first studied neutrophil shape changes adhered to E-selectin, ICAM-1 and CXCL8 coated flow chambers under flow conditions. Human neutrophils were introduced into the flow chambers and changes in cell shape was monitored over time (Fig. 5A). sPIF treated cells remained rounder, did not polarize and form cell protrusions as reflected by significant lower cell perimeter than controls (Fig. 5B). Next, we subjected the cells to increasing shear forces after allowing them to settle on the substrate for 3min in the flow devices. sPIF treated neutrophils detached at lower shear rates compared to control cells (Fig. 5C), indicating that PIF mediated impairment of cytoskeletal rearrangements prevents adhesion strengthening *in vitro*. To test the inability of PIF treated neutrophils to firmly adhere under flow conditions *in vivo*, we performed IVM of unstimulated cremaster muscles (trauma model) and analyzed neutrophil adhesion to postcapillary venules before and after intra-arterial injection of CXCL1 (Fig. 5D). Application of CXCL1 to *WT* mice pretreated with scrPIF induced a significant increase of adherent neutrophils within 30s that did not decrease over 5min of observation (Fig. 5E). sPIF treated mice also exhibited a significant increase in adherent neutrophils 30s post CXCL1 injection, but this number subsequently decreased following 5min of observation. This indicated that inhibition of K_V_1.3 by PIF increases susceptibility to physiological shear forces *in vivo*. In addition, we analyzed induction of neutrophil adhesion in these experiments (number of adherent cells mm^−2^ 30s post injection relative to before), yet we did not detect any differences in relative adhesion 30s post injection (Supplemental Fig. S4A). Based on these observations, it appeared that integrin inside-out signaling is not affected by PIF activity. To study β_2_ integrin activation in more detail, we monitored CXCL8 induced changes in β_2_ integrin conformation in human neutrophils using antibodies that recognize specific integrin affinity states (clone KIM127: intermediate affinity; clone mAB24: intermediate and high affinity) ^33^ and flow cytometry. Pre-incubation with sPIF did not affect the binding capacity of either antibody compared to controls (Fig.5F-I), further demonstrating that PIF does not impair GPCR-mediated inside-out signaling of a shift from low to high affinity conformation. Of note, quantitative surface expression of the β_2_ integrins LFA-1 and MAC-1 were not affected by sPIF (Supplemental Fig. S4B-D). Moreover, the binding capacity of murine neutrophils to ICAM-1 upon CXCL1 stimulation was not affected by the presence of sPIF either (Supplemental Fig. S4E,F). These results suggest that PIF impairs neutrophil adhesion *in vivo* and *in vitro* by reducing adhesion strengthening via β_2_ integrin outside-in signaling, mediated by interfering with K_V_1.3 activity. To support this, we performed IVM of TNF-stimulated post capillary venules in the mouse cremaster muscle using *WT* mice treated with a combination of sPIF and the K_V_1.3 inhibitor PAP-1 as well as K_V_1.3-deficient mice (*Kcna3^−/-^*) treated with sPIF. Neither concomitant administration of sPIF and PAP-1 together, nor sPIF injection into *Kcna3^−/-^* mice did further reduce the number of adherent neutrophils compared to sPIF treatment alone or scrPIF+PAP-1 treatment (Fig. 5J, Supplemental Table S4). Taken together, these data indicate that PIF exerts its function on neutrophil arrest and adhesion strengthening by blocking K_V_1.3.

**Figure 5.**
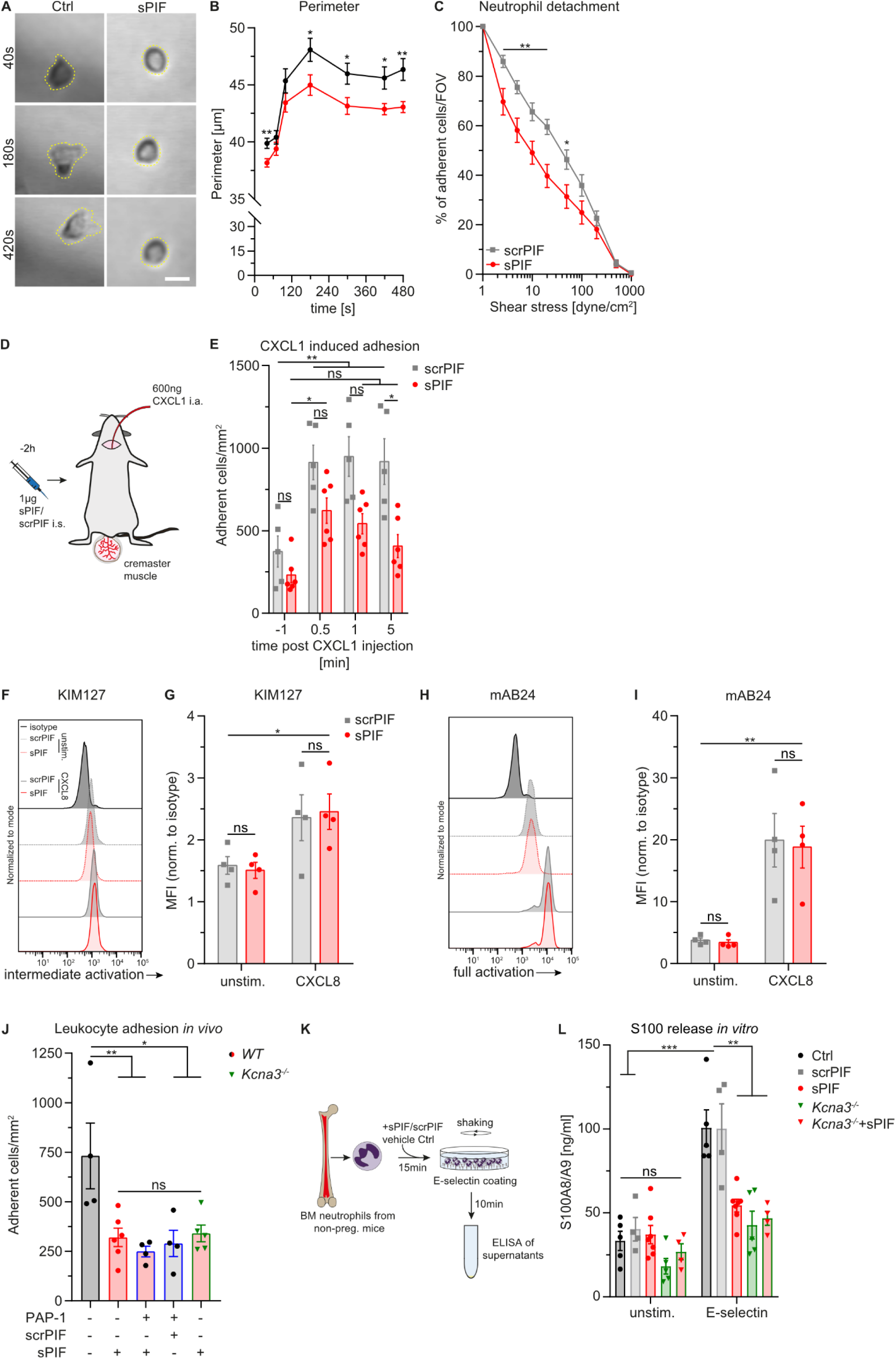
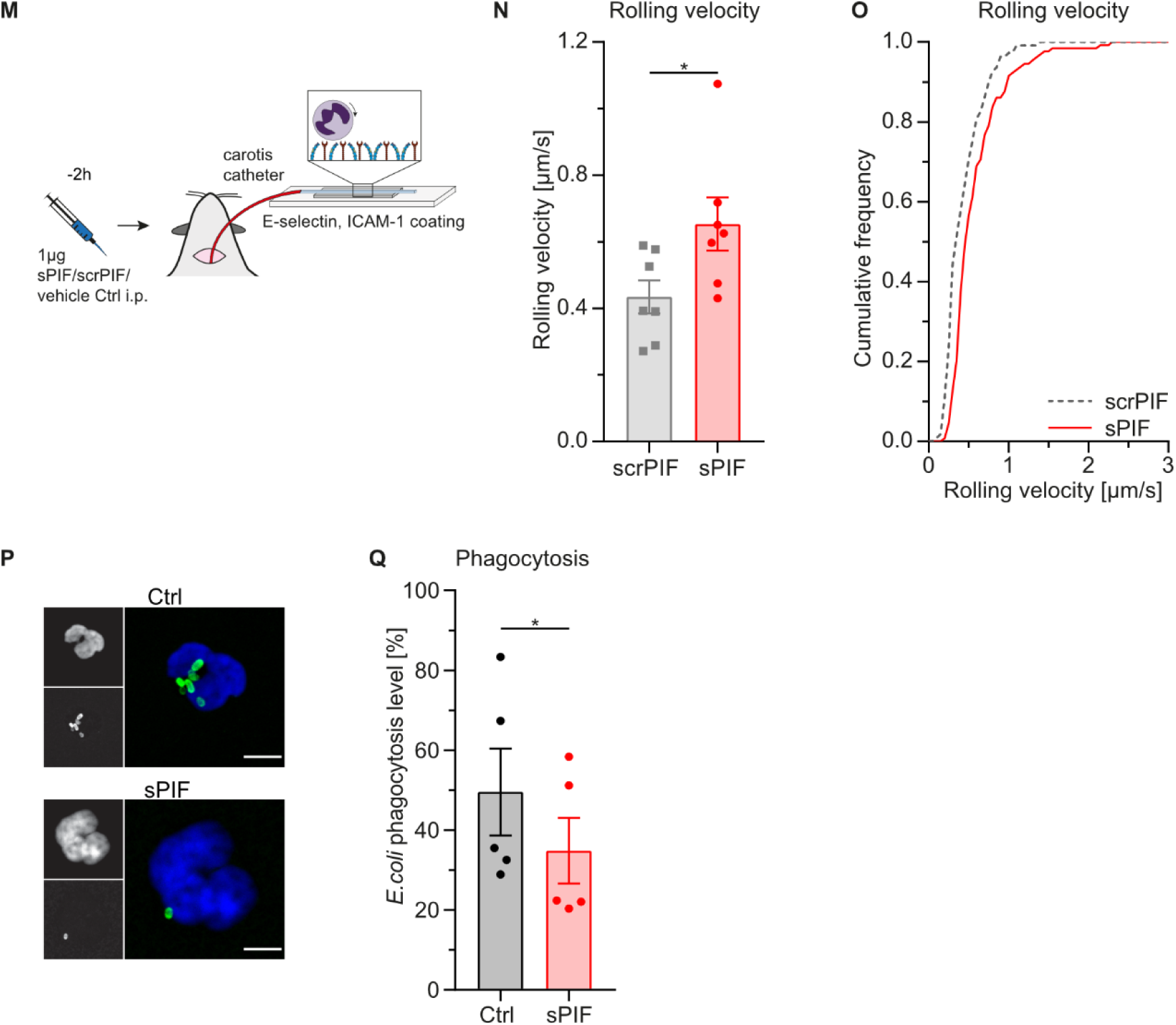
PIF-mediated inhibition of K_V_1.3 impairs post-arrest modifications, S100A8/A9 release and phagocytosis in neutrophils. (**A**) Human neutrophils pretreated with 300nM sPIF or vehicle (Ctrl) were introduced into E-selectin/ICAM-1/CXCL8 coated flow chambers and monitored over time (representative micrographs of neutrophils 40s, 180s and 420s after attachment, scale bar: 10µm). (**B**) Cell perimeters were analyzed over time (52 (Ctrl) and 40 (sPIF) cells from n=5 independent experiments, repeated unpaired Student’s t-tests). (**C**) sPIF and scrPIF (300nM) pretreated human neutrophils were exposed to stepwise increasing shear rates in E-selectin/ICAM-1/CXCL8-coated flow chambers and percentage of remaining neutrophils FOV^−1^ was quantified after each step (n=15-18 flow chambers of 8-9 independent experiments, repeated unpaired Student’s t-tests). (**D**) Schematic of experimental design of CXCL1 application during IVM of the mouse cremaster muscle. (**E**) Neutrophil adhesion was quantified before and after i.a. injection of 600ng CXCL1 in *WT* mice pretreated with 1µg sPIF or scrPIF (n≥5mice per group, 2-way ANOVA, Tukey’s multiple comparison). (**F-I**) β_2_ integrin activation (KIM127: intermediate activation; mAB24: full activation) was assessed in isolated human neutrophils pretreated with 300nM sPIF or scrPIF and stimulated with 10nM CXCL8 and analyzed by flow cytometry (**F,H**: representative flow cytometry blots, MFI: median fluorescence, n=4 independent experiments, 2-way RM ANOVA, Sidak’s multiple comparison). (**J**) Neutrophil adhesion to postcapillary venules in the cremaster muscle of WT and *Kcna3^−/-^* i.s. injected with sPIF, scrPIF (both 1µg), PAP-1 (90µg) or vehicle 3h and stimulated i.s. with TNF 2h prior to IVM (n≥4 mice per group; 1-way ANOVA, Tukey’s multiple comparison). (**K**) Experimental design of E-selectin-triggered S100A8/A9 release assays *in vitro*. (**L**) S100A8/A9 levels of PBS (unstim.) or E-selectin stimulated bone marrow neutrophils from WT and *Kcna3^−/-^* mice pretreated with sPIF, scrPIF (both 300nM) or vehicle control for 15min (n≥4mice per group; 2-way RM ANOVA; Sidak’s multiple comparison). (**M**) Schematic of *ex vivo* flow chamber assays. (**N**) Mean rolling velocities and (**O**) cumulative distribution of rolling velocities of neutrophils from WT mice i.p. injected with sPIF or scrPIF (both 1µg) and perfused through E-selectin and ICAM-1 coated flow chambers (n=7 mice per group; unpaired Student’s t-test; cumulative frequency: n= 109 (scrPIF) and n= 129 (sPIF) cells). (**P**) Phagocytic activity of neutrophils was assessed using fluorescent *E. coli* particles in human whole blood pre-incubated with sPIF (300nM) or vehicle (Ctrl) (representative micrographs, scale bar =5µm) and (**Q**) analyzed by flow cytometry (n=5 independent experiments, paired Student’s t-test). Data is presented as representative micrographs, mean±SEM, representative flow cytometry plots and as cumulative frequency.

Besides its role in the regulation of adhesion and post-arrest modifications, we recently revealed that potassium efflux via K_V_1.3 is required for NLRP3-dependent S100A8/A9 release upon engagement with E-selectin ^29^. S100A8/A9 is the most abundant cytosolic protein in neutrophils and functions as an alarmin or danger associate molecular pattern (DAMP) ^34^. We therefore asked whether inhibition of K_V_1.3 by PIF impairs E-selectin-triggered S100A8/A9 release as well. To address this, we stimulated WT and *Kcna3^−/-^* neutrophils pretreated with sPIF, scrPIF or vehicle control for 10min with E-selectin and measured S100A8/A9 concentration in the supernatant by ELISA (Fig. 5K). E-selectin stimulation increased S100A8/A9 levels in vehicle or scrPIF treated WT neutrophils whereas no significant change was observed in the supernatants of WT neutrophils preincubated with sPIF (Fig. 5L). Of note, S100A8/A9 levels were similar in control treated and PIF treated neutrophils from *Kcna3^−/-^*mice, demonstrating that PIF-mediated impairment of S100A8/9 release is predominantly mediated by inhibition of K_V_1.3. Released S1008/A9 binds to TLR4 on rolling neutrophils in an autocrine fashion resulting in an activation of β_2_ integrins and a deceleration of rolling neutrophils ^29, 35, 36^. To investigate whether impaired S100A8/A9 release in the presence of PIF alters rolling velocities, we performed *ex vivo* flow assays, perfusing E selectin and ICAM-1-coated flow chambers with whole blood from WT mice that were injected with sPIF or scrPIF 2h prior to the experiment (Fig. 5M). In line with the S100A8/A9 release data, neutrophils from sPIF treated animals exhibited significant increased rolling velocities on E-selectin and ICAM-1-coated flow chambers compared to scrPIF injection (Fig. 5N, O). Further, isolated human neutrophils pretreated with sPIF resulted in significant increased rolling velocities in E-selectin and ICAM-1 coated flow chambers compared to scrPIF control treatment too (Supplemental Fig. S5A-C), demonstrating that reduced S100A8/A9 release is associated with impaired slow neutrophil rolling *ex vivo* and *in vitro*.

K_V_1.3 regulated sustained Ca^2+^ influx is a prerequisite for sufficient phagocytosis ^28^ and neutrophils from pregnant women exhibit a reduced phagocytic capacity ^37^ which was linked to a serum factor ^38^. Accordingly, we hypothesized that PIF is a serum factor impairing phagocytosis during pregnancy. To assess this, we incubated human whole blood with fluorescent *E. coli* particles and analyzed phagocytic activity by confocal microscopy and flow cytometry. Pre-treatment with sPIF resulted in reduced phagocytosis of *E.coli* particles compared to vehicle treatment (Fig. 5P,Q; Supplemental Fig. S5D-F), demonstrating that PIF reduces phagocytic capacity of neutrophils.

In conclusion, we demonstrate that PIF acts as an endogenous inhibitor of K_V_1.3, thereby interfering with K_V_1.3-regulated neutrophil functions, including post-arrest modification/adhesion, active alarmin release and phagocytosis.

## Discussion

Local immune cell regulation in close proximity to the fetus is a central prerequisite for successful pregnancy and dysregulation of intrauterine immune cell composition is associated with severe pregnancy-related complications ^39^. How systemic immunity is regulated in the becoming mother is less clear. Although pregnant women exhibit elevated WBC counts throughout gestation, which is mainly due to an increased number of circulating neutrophils ^40, 41^, they are at the same time more susceptible to certain infections ^24, 42^, indicating reduced cellular immunity during pregnancy. In the present study we mechanistically advance the knowledge on functional neutrophil modulation in this critical life period by showing that pregnancy is associated with altered neutrophil recruitment *in vivo* and *in vitro*, both in mice and humans.

By using synthetic protein and blocking antibodies, we were able to mechanistically link altered neutrophil functions during pregnancy to the presence of the trophoblast-derived small peptide PIF in the maternal serum. In addition, exogenous PIF interferes with neutrophil recruitment in activated cremaster muscles as well as in *Legionella* pneumonia. These observations reflect impaired neutrophil post-arrest modifications, reduced S100A8/A9 release, and altered phagocytic activity via inhibition of the voltage-gated potassium channel K_V_1.3 on neutrophils. An earlier study suggested that PIF might interact with K_V_1.3 ^27^. Using an electrophysiological approach, we provide clear evidence that PIF is an endogenous inhibitor of K_V_1.3, blocking its activity with an IC_50_ of 10±5nM. PIF concentrations of around 50-60nM as reported in sera from pregnant women ^14^ are therefore very likely to sufficiently block K_V_1.3 on circulating neutrophils. Many neutrophil functions are regulated by changes in [Ca^2+^]_i_ including cytoskeletal rearrangement, integrin function, phagocytosis, and ROS production ^43^. Recently, we have shown that neutrophil expressed K_V_1.3 ensures long lasting Ca^2+^ signaling by stabilizing membrane potential ^28^. Genetic deletion or pharmacological inhibition of K_V_1.3 in neutrophils resulted in disturbed cytoskeletal rearrangement under flow conditions (integrin outside-in signaling) and an increased susceptibility to physiological shear forces. This is in line with our findings that PIF impairs sustained Ca^2+^ signaling and leading to disturbed cell spreading *in vitro* and neutrophil adhesion *in vitro* and *in vivo*. Interestingly, neutrophils exhibit altered Ca^2+^ oscillations upon contact with trophoblast cells *in vitro* which was attributed to changes in glucose metabolism ^44^. Our observations that trophoblast-derived PIF alters [Ca^2+^]_i_ via K_V_1.3 inhibition in neutrophils adds an additional explanation contributing to the observed phenotype.

Besides its role in sustaining Ca^2+^ entry during neutrophil activation and in regulating post-arrest modifications/arrest and phagocytosis, we have recently linked K_V_1.3 activity to S100A8/A9 release from neutrophils ^29, 45, 46^. K^+^ efflux via K_V_1.3 is required for E-selectin-mediated rapid NLRP3 inflammasome activation and ensuing GSDMD pore formation by which S100A8/A9 is secreted. S100A8/A9 in turn can bind in to TLR4 and activate β_2_ integrins in an autocrine manner, triggering neutrophil adhesion ^35^. In this context, PIF application prevented S100A8/A9 release *in vitro*, resulting in concomitant increased rolling velocities.

The fact that K_V_1.3 is expressed on many immune cells ^47^ regulating among other things cell proliferation and cytokine production ^48^ indicates that PIF does not only modulate neutrophil functions, but might be a central modulator of local and systemic immunity during pregnancy. In fact, immune-modulatory properties of PIF in general have been documented in mouse models outside pregnancy by showing that continuous PIF administration reduces the severity of multiple sclerosis (MS) and type I diabetes mellitus (TIDM), accompanied by reduced immune cell infiltration into affected tissues ^17, 18^. Further, PIF application diminished Graft-versus-Host disease after bone marrow transplantation in mice ^49^. In addition to these observations, PIF was shown to create a pro-receptive milieu in the decidua following conception ^27^ and during trophoblast invasion ^12, 50^. Further, PIF enhances local progesterone activity, increases steroid secretion and enhances the expression of pro-tolerogenic HLA molecules in cytotrophoblasts ^51^, highlighting its important role during pregnancy.

In summary, we demonstrate that PIF suppresses neutrophil functions during pregnancy via its direct effect on the voltage-gated potassium channel K_V_1.3. Inhibition of K_V_1.3 by PIF impairs intracellular Ca^2+^ signaling and reduces neutrophil recruitment into inflamed tissue as well as neutrophil phagocytosis and NLRP3 inflammasome-dependent S100A8/A9 release. This identifies PIF as a critical and specific factor that modulates neutrophil effector functions during pregnancy. These findings may open a broad range of new diagnostic and therapeutic opportunities related to PIF during pregnancy and more broadly in treatment of inflammation.

## Material and Methods

### Study approval

Animal experiments were approved by the Government of Oberbayern (AZ 55.2-1-54-2531-122/12, - 229/15, ROB-55.2-2532.Vet_02-18-22) and the Government of Baden, Germany (AZ G21/025). Blood sampling from healthy volunteers was approved by the ethic committee from the Ludwig-Maximilians-Universität München, Munich, Germany (AZ 22-0534 und 611-15) and by the institutional review board protocol at the University of California, Davis, USA (#235586-9).

### Mice

*Catchup^IVM-red^* (*Ly6g^Cre-tom/wt^, ROSA26^Rtom/wt^, C57BL/6*) mice ^25^ were kindly provided by M. Gunzer, University Duisburg-Essen, Germany. *Kcna3^−/-^*(*Kcna3^tm1Lys^*) mice ^52^ were purchased from Jackson Laboratories and maintained on a C57BL/6NCrl background. C57BL/6NCrl (*WT*) mice were obtained from Charles River Laboratories (Sulzfeld, Germany). All mice were maintained at the Walter Brendel Center for Experimental Medicine, LMU, Munich, at the Biomedical Center, LMU, Planegg-Martinsried, and at Center for Experimental Models and Transgenic Service, University of Freiburg, Germany. 8-25 weeks old male and female mice were used for all experiments. For *in vivo* experiments, mice were anaesthetized via i.p. injection of a combination of ketamine/xylazine (125mg kg^−1^ and 12.5mg kg^−1^ body weight, respectively in a volume of 0.1ml 8g^−1^ body weight). Adequacy of anesthesia was ensured by testing the metatarsal reflex, monitored throughout the experiment and refreshed once an hour (0.03ml 8g^−1^ body weight). All mice were sacrificed by cervical dislocation.

### Neutrophil isolation

Mouse: Bone marrow neutrophils were isolated either by density centrifugation using Percoll® gradient (Sigma-Aldrich) or by negative selection (Neutrophil enrichment kit, STEMCELL Technologies). Human: Human neutrophils were isolated from heparinized whole blood samples of healthy volunteers by density centrifugation using Polymorphprep (AXIS-SHIELD PoC AS) or a direct negative selection kit (STEMCELL Technologies).

Cells were resuspended in Hank’s balanced salt solution (HBSS) buffer [containing 0.1% of glucose, 1mM CaCl_2_, 1mM MgCl_2_, 0.25% BSA, and 10mM HEPES (Sigma-Aldrich), pH7.4].

### Intravital microscopy

Intravital microscopy (IVM) of the mouse mesentery was carried out in pregnant (day E13.5) and non-pregnant female *Catchup^IVM-red^*mice that received an intra peritoneal (i.p.) injection of 1µg recombinant murine (rm)TNF (R&D Systems) 3h prior to investigation. The carotid artery of anesthetized mice was cannulated for later blood sampling (using a ProCyte Dx; IDEXX Laboratories) and administration of fluorescent beads (Flouresbrite® YG Microspheres, 1µm) for later offline determination of blood flow velocities. Mesenteric vessels were exposed and constantly superfused with thermo-controlled bicarbonate buffer ^53^. IVM was conducted on a Zeiss Axio Examiner.D1 microscope equipped with a 40x objective (0.75 NA, water immersion), an Axiocam 702 CMOS camera and a Colibri epifluorescence LED excitation light source (all Zeiss). Mesenteric venules were recorded using Zen software (Zeiss) for later analysis. Neutrophil rolling flux, adhesion, and rolling velocities as well as vessel diameters, vessel lengths, and blood flow velocities were assessed offline using FIJI software ^54^ including the MTrackJ plugin ^55^ for determination of rolling and blood flow velocities.

IVM of the mouse cremaster muscle was performed as previously described ^56^. Briefly, *WT* or *Kcna3^−/-^*mice received an intrascrotal (i.s.) injection of either synthetic (s)PIF (1µg/mouse; BioSynthesis and Bachem), scrambled PIF (scrPIF; sequence: GRVDPSNKSMPKDIA; 1µg/mouse; BioSynthesis), 5-(4-Phenoxybutoxy)psoralen (PAP-1; 90µg/mouse; Sigma-Aldrich), vehicle (Ctrl; 0.25%DMSO/PBS) or a combination of the mentioned substances as indicated 1h prior to rmTNF stimulation (500ng i.s.). 2h after induction of inflammation, the carotid artery of anesthetized mice was cannulated for later blood sampling and the cremaster muscle was dissected and constantly superfused with thermo-controlled bicarbonate buffer. IVM was conducted on an OlympusBX51 WI microscope, equipped with a 40x objective (Olympus, 0.8NA, water immersion objective) and a CCD camera (Kappa CF 8 HS). Postcapillary venules were recorded using VirtualDub software for later analysis (see above). Centerline velocity of each venule was measured with a dual photodiode (Circusoft Instrumentation). After IVM, cremaster muscles were removed, fixed with 4% paraformaldehyde (PFA), and stained with Giemsa (Merck) for quantification of perivascular neutrophils. Analysis was carried out on a Leica DM2500 microscope, equipped with a 100x objective (Leica, 1.4NA, oil immersion) and a Leica DMC2900 CMOS camera.

For CXCL1 induced adhesion experiments, *WT* mice received an i.s. injection of 1µg sPIF or scrPIF, respectively 2h prior to IVM. Leukocyte adhesion was analyzed before and after application of rmCXCL1 via the carotid catheter (600ng, Peprotech) in one postcapillary venule per mouse.

### *Legionella pneumophilia* infection

Infection with *Legionella pneumophilia* (*L.p.*) was carried out using the strain JR32 as previously described ^57^. Briefly, bacteria were plated on buffered charcoal yeast extract (BCYE) agar (Thermo Fisher). The optical density (OD600) of one and more single colonies was measured using an optical spectrophotometer. Subsequently, the suspension was plated on BCYE agar plates to create a standard curve. For infection, single colonies were picked, suspended in sterile PBS and the OD600 was measured to determine and adjust the infectious dose of approximately 5×10^6^ CFU.

To extract respiratory immune cells, *WT* mice were sacrificed via cervical dislocation and subsequently perfused with PBS through the right cardiac ventricle. Next, the right lung (cranial, caudal, middle and accessory lobes) was homogenized and incubated in PBS containing 10% FCS, Hyaluronidase (1mg ml^−1^, Sigma Aldrich), Collagenase IV (0.25mg ml^−1^, Worthington) and DNase I (0.25mg ml^−1^, Sigma Aldrich) for 1h at 180rpm, 37°C. The obtained cell suspension was filtered with a 70µm cell strainer and remaining erythrocytes were lysed with 1x RBC lysis buffer (eBioscience) followed by washing. Cells were then incubated with anti-CD16/32 antibody (clone 93) for 10min on ice, diluted with 1% FBS, 2mM EDTA in PBS (FACS buffer), and stained against CD45 (30-F11, eF450 conjugated), CD11b (clone M1/70, PE-Cy7 conjugated), Ly6C (clone HK1.4, BV605 conjugated) and Ly6G (clone 1A8, APC conjugated) for 30min at 4°C in FACS buffer. Finally, dead cells were stained with fixable viability dye eFluor780 (eBioscience) according to manufacturer’s instruction. Samples were analyzed with a 4-laser flow cytometer (Cytoflex S, Beckman Coulter) and data were processed with the Kaluza software (v2.3, Beckman Coulter).

### Surface expression of rolling- and adhesion-relevant molecules

Peripheral blood from anesthetized pregnant and non-pregnant WT mice and WT mice pretreated with 1µg sPIF or vehicle control i.s. for 2h was obtained by retro-orbital puncture. Whole blood samples were stained against: CD11a (clone M17/4; APC conjugated), CD11b (clone M1/70; BV510 conjugated), CD18 (clone C7/16; FITC conjugated), CD62L (clone MEL-14; FITC conjugated), CD162 (clone 2PH1; PE conjugated), CXCR2 (clone 242216; APC conjugated), and CD44 (clone IM7; BV570 conjugated), all 5µg ml^−1^. Cells were fixed and erythrocytes lysed using FACS-lysing solution (BD Bioscience). Samples were analyzed using a CytoFlexS flow cytometer (Beckmann Coulter) and FlowJo software. Ly6G^+^ cells (clone 1A8, PB conjugated; BioLegend) were defined as neutrophils.

### Murine flow chamber assays

Flow chamber assays were performed as previously described ^28^. Rectangular borosilicate glass capillaries (40×400µm; VitroCom) were coated with rmE-selectin (CD62E Fc chimera; 20µg mL^−1^), rmICAM-1 (ICAM-1 Fc chimera; 15µg mL^−1^; both R&D Systems), and rmCXCL1 (15µg mL^−1^) for 3h at room temperature (RT), subsequently blocked over night (ON) with PBS containing 5% casein and washed with PBS. For serum experiments, 1×10^5^ isolated bone marrow neutrophils from non-pregnant *WT* mice (female and male; in 10µl HBSS) were incubated with serum from pregnant or non-pregnant mice (total volume of 50µl) for 15min at RT and subsequently introduced into the flow chambers using a high-precision syringe pump (Harvard Apparatus). For PIF blockade, cells were additionally incubated with anti-PIF antibodies (clone 852/F2, InVivo BioTech Services) or mouse IgG1,κ isotype control (clone MOPC-21 Biolegend), both 60µg ml^−1^ and rPIF as indicated. Experiments were conducted on a Zeiss Axio Examiner.D1 microscope equipped with a 20x objective (1.0 NA, water immersion) and an Axiocam 702 CMOS camera and recorded using Zen software (Zeiss).

For whole blood experiments, whole blood was collected from anesthetized pregnant and non-pregnant *WT* mice via the carotid catheter, heparinized and incubated with either 300nM sPIF or vehicle control (Ctrl) for 15min at RT. Incubated blood was perfused through the flow chambers at a constant shear rate level of 2.7dynes cm^−2^ using a high-precision syringe pump. Movies were recorded with an OlympusBX51 WI microscope, equipped with a 20x objective (Olympus, 0.95NA, water immersion objective) and a CCD camera (Kappa CF 8 HS) and VirtualDub software. Neutrophil rolling, rolling velocities and adhesion were quantified offline with the help of FIJI software.

For *ex vivo* flow chambers assays, WT mice were injected i.p. with sPIF or scrPIF (1µg) and 2h later connected via a carotid catheter to rmE-selectin (20µg mL^−1^) and rmICAM-1 (15µg mL^−1^) coated flow chambers recorded for later offline analysis of rolling velocities.

### Patch clamp of K_V_1.3 overexpressing HEK-293 cells and isolated human neutrophils

HEK-293 cells were cultured in DMEM supplemented with 10% FCS and 1% penicillin/streptomycin at 37°C in humidified atmosphere (5% CO_2_). Cells were transiently co-transfected with cDNA plasmid endcoding human K_V_1.3 (kindly provided by Prof. Dr. Grissmer, University of Ulm) and Vivid Colors™ pcDNA™6.2/C-EmGFP-DEST Vector (Invitrogen) using Lipofectamine 3000 (Invitrogen). Cells were used the day after transfection.

Isolated human neutrophils were seeded on poly-D-lysine coated coverslips. Cells were subjected to patch-clamp experiments in whole-cell configuration as follows: cells were clamped at a holding potential of −80mV intermitted by repeated 200ms voltage steps from −80mV to +40mV using a 10mV interval applied every 30s. Current maxima at +40mV were used for the calculation of K_v_1.3 current amplitudes. Currents were normalized to cell size as current densities in pA/pF. Capacitance was measured using the automated capacitance cancellation function of the EPC-10 (HEKA, Harvard Bioscience). Patch pipettes were made of borosilicate glass (Science Products) and had resistance of 2–3.5MΩ. K_v_1.3 currents were inhibited with 300nM and varying concentrations of sPIF, 10nM PAP-1 (5-(4-Phenoxybutoxy)psoralen; Sigma-Aldrich). Standard extracellular solution contained: 140mM NaCl, 2.8mM KCl, 2mM MgCl_2_, 1mM CaCl_2_, 10mM HEPES, 11mM glucose (pH 7.2, 300mOsm). Intracellular solution contained: 134mM KF, 2mM MgCl_2_, 1mM CaCl_2_, 10mM HEPES, 10mM EGTA (pH 7.2, 300mOsm). Solutions were adjusted to 300mOsm using a Vapro 5520 osmometer (Wescor Inc). Channel blocker were either added to the bath solution at least 15min prior to electrophysiological recordings or directly applied via an application pipette using constant pressure. To determine IC_50_ values for inhibitory effects of PIF on K_v_1.3 currents, data were fitted with the following equation: *E(c) =E_min_+(E_max_−E_min_) x (1/(1 + (IC_50_/c)^h^))*; with *E* being the effect (current inhibition) at a given concentration *c* of inhibitor, *E_min_* the minimal effect (current inhibition), *E_max_* the maximally achievable effect, *IC_50_* the half-maximal concentration and *h* the Hill factor.

### Calcium imaging of human neutrophils

Isolated human neutrophils were loaded with Rhod-2 AM (1µM, ex/em: 552/581) with or without Thapsigargin (2µM, both Thermo Fischer Scientific) for 20min at RT in the dark and subsequently incubated with 300nM sPIF or scrPIF, respectively for 15min. Cells were placed into 48-well plates and allowed to settle for 60s prior to addition of Ca^2+^ (1.5mM) or 10nM rhCXCL8 (Shenandoah Biotechnology), respectively. Images were taken (1 fps) using a Nikon Eclipse TE2000-S microscope (20x phase contrast air objective; 0.45NA) equipped with a 16-bit digital CMOS camera (Andor ZYLA) with NIS Elements imaging software. All images were analyzed using FIJI Software.

### Human flow chamber assays

The effect of pregnant serum on neutrophil adhesion was studied in µ-slides VI^0.1^ (ibidi) flow chambers coated with rhE-selectin (CD62E Fc chimera; 5µg ml^−1^; R&D Systems), rhICAM-1 (4µg ml^−1^; R&D Systems) and CXCL8 (10µg ml^−1^; Peprotech). 5×10^5^ isolated neutrophils from non-pregnant female or male donors were incubated with serum from pregnant (week 20-25 of gestation) and non-pregnant donors (males) for 15min at 37°C, introduced into the flow devices and after 3min superfused with respective serum at a constant shear stress level of 2dyne cm^−2^ using a high-precision syringe pump (Harvard Apparatus). Experiments were conducted on a Zeiss Axio Examiner.D1 microscope equipped with a 20x objective (1.0 NA, water immersion) and an Axiocam 702 CMOS camera and recorded using Zen software (Zeiss).

Neutrophil spreading was studied in rhE-selectin, rhICAM-1 and CXCL8 coated rectangular borosilicate glass capillaries (0.2×2.0mm; CM Scientific). Flow chambers were perfused with isolated human neutrophils pretreated with 300nM sPIF or vehicle control for 15min and hFc-block (human TruStain FcX; BioLegend) for 5min at RT at constant shear stress level of 1dyne cm^−2^ using a high-precision pump. Cell shape changes were recorded with a Zeiss Axioskop2 (equipped with a 20x water objective, 0.5NA and a Hitachi KP-M1AP camera), VirtualDub software and analyzed using FIJI software.

To assess neutrophil detachment to increasing shear forces, isolated human neutrophils were incubated with 300nM sPIF or scrPIF for 15min and hFc-block for 5min at RT and subsequently introduced into in rhE-selectin, rhICAM-1 and CXCL8 coated µ-slides VI^0.1^ (ibidi) flow chambers. Cells were allowed to settle for 3min until 1dyne cm^−2^ was applied for 1min to remove non-attached cells. Flow rates were then increased every 30s and the fraction of attached neutrophils was counted at each time interval. Experiments were carried out on a ZEISS AXIOVERT 200 microscope, equipped with a ZEISS 10x objective NA0.25 and a SPOT RT ST camera (Diagnostic Instruments) and MetaMorph software.

Analysis of rolling velocities was carried out in rhE-selectin (5µg ml^−1^) and rhICAM-1 (4µg ml^−1^) coated flow chambers perfused with isolated human neutrophils at constant shear stress level of 1dyne cm^−2^. Cells were pretreated with hFc-block for 5min at RT and subsequent incubation with 300nM sPIF or scrPIF, respectively for 15min.

### Integrin activation and soluble ICAM-1 binding assay

Integrin activation was carried out as previously described ^28^. Isolated human neutrophils were incubated for 15min at RT with 300nM sPIF or scrPIF, respectively, and subsequently activated with 10nM CXCL8 in the presence of anti-CD18 antibodies (clone KIM127, recognizing intermediate activated CD18, INVIVO; clone mAB24, recognizing fully activated CD18, Hycult Biotech; both 10µg ml^−1^) for 5min at 37°C. Stimulated was stopped by adding ice-cold FACS lysing solution (BD BioScience). Secondary PE-conjugated anti-mouse antibody (0.5µg ml^−1^, Pharmingen) was used to quantify binding capacity of anti-CD18 antibodies. Neutrophils were defined as CD15^+^/CD66^+^ population (clones W6D3, FITC-conjugated and G10M5, APC-conjugated; both 5µg ml^−1^, BioLegend). In addition to CD18, Overall surface expression of CD11a (clone HI111) and CD11b (clone ICRF44, both 5µg ml^−1^, BioLegend) were assessed.

Bone marrow neutrophils from *WT* mice pre-treated with 300nM sPIF or scrPIF, respectively were stimulated for 3min at 37°C with 10nM CXCL1 in the presence of soluble rmICAM-1 (40µg ml^−1^, R&D Systems, with human IgG1 tag), pre-complexed with biotin-conjugated anti-human Fc (10µg ml^−1^, eBioscience) and PerCP-Cy5.5-conjugated streptavidin (2µg ml^−1^, BioLegend). Stimulation was stopped by ice-cold FACS lysing solution. Neutrophils were defined as Ly6G^+^ population (clone 1A8, PB-conjugated, 5µg ml^−1^, BioLegend). Samples were analyzed by flow cytometry (Beckman Coulter Gallios) and FlowJo software.

### Phagocytosis

Heparinized human whole blood was pre-treated with 300nM sPIF or vehicle control for 15min at RT and subsequently incubated with *Escherichia coli* particles (pHrodo Green *E. coli* BioParticle Phagocytosis Kit, Thermo Fisher Scientific) for 30min at 37°C (or at 4°C as negative control). Phagocytosis was stopped and cells were fixed using the provided fixative according to the manufacturer’s protocol (including RBC lysis). Samples were analyzed by flow cytometry using a Beckman Coulter Gallios flow cytometer and FlowJo software. Neutrophils were defined as CD15^+^/CD66b^+^ (clone W6D3; APC-conjugated; clone G10F5, PB-conjugated; both 5µg ml^−1^, BioLegend) positive population.

For representative confocal micrographs, phagocytosis was stopped by transferring samples on ice. Neutrophils were isolated by direct negative selection and seeded on poly-L-lysine (0.1%; Sigma-Aldrich) coated object slides (Ibidi). Cells were fixed (2% PFA), stained with DAPI (Invitrogen) for 5min at RT and embedded with ProLong Diamond Antifade mounting medium. Images were acquired using a SP8X WLL microscope, equipped with a HC PL APO 40x/1.30NA oil immersion objective. Images were processed (including removal of outliers and background subtraction) using FIJI software.

### S100A8/S100A9 release *in vitro*

E-selectin-induced S100 release from isolated bone marrow neutrophils was carried out as reported elsewhere ^35^. Briefly, 24 well plates (Suspension cell plates, Greiner Bio-One) were coated with PBS/0.1% BSA or rmE-selectin (10µg mL^−1^), respectively at 4°C ON, blocked with 5% casein for 3h at RT and subsequently washed with PBS. Isolated bone marrow neutrophils from *WT* and *Kcna3^−/-^* mice were pretreated with 300nM sPIF, scrPIF (both 300nM) or vehicle control for 15min at 37°C with 5% CO_2_. 5×10^5^ cells in 500µl were added to the 24 wells and incubated for 10min at 37°C with 5% CO_2_ under shaking conditions. Sell supernatants were analyzed by ELISA to determine S100A8/S100A9 concentrations.

### Statistics

Data are presented as mean±SEM, cumulative frequency, representative images/traces/histograms as detailed in the figure legends. Group sizes were chosen based on previous experiments. GraphPad Prism software (GraphPad software) and Adobe Illustrator were used to analyzed data and illustrate graphs. Statistical testing was carried out according to the number of experimental groups. For pairwise comparisons, an unpaired und paired Student’s t-test, respectively was used. For more than groups, one-way or two-way ANOVA with Tukey’s or Sidak’s multiple comparison post hoc tests were performed. Data distribution was assumed to be normal, but this not formally tested. P values of <0.05 were considered statistically significant and indicated as follows: ^∗^p < 0.05; ∗∗p < 0.01; ∗∗∗p < 0.001.

## Acknowledgements

This work was supported by the German Research Foundation (DFG) collaborative research grants - TRR359 – project ID 491676693 (B2 to RI and MS, A02 to PH, Z01 to MB, LAUNCH1 seed funding to RI and SB), TRR332 – project ID 449437943 (to TV, MS). PH received further funding by the DFG TRR167 (Project ID 259373024) and CRC1160 (Project ID 256073931). We thank Dorothee Gössel for excellent technical assistance, the core facilities Bioimaging and animal models at the Biomedical Center (LMU Munich) for their support and M. Gunzer (University of Duisburg-Essen) and S. Grissmer (University of Ulm) for providing critical reagents. In addition, we thank all volunteer blood donors.

## Author contribution

RI designed and conducted experiments, analyzed data and wrote the manuscript. WN, JMF, AB, SB, VAM, AK, MB, IF, ECMI, JBMR, MP, LMW, MN, IR, AY, MR, and TV acquired and analyzed data. MK, MM, EB, MB, and CK provided their expertise and critical reagents. SIS, PH, AI, and SZ designed experiments, analyzed results and provided their expertise. MS designed experiments and wrote the manuscript.

