## Supplemental Figures for "Preimplantation factor (PIF) is an endogenous inhibitor of potassium channel K_V_1.3 regulating neutrophil function during pregnancy"

Supplemental Figure S1

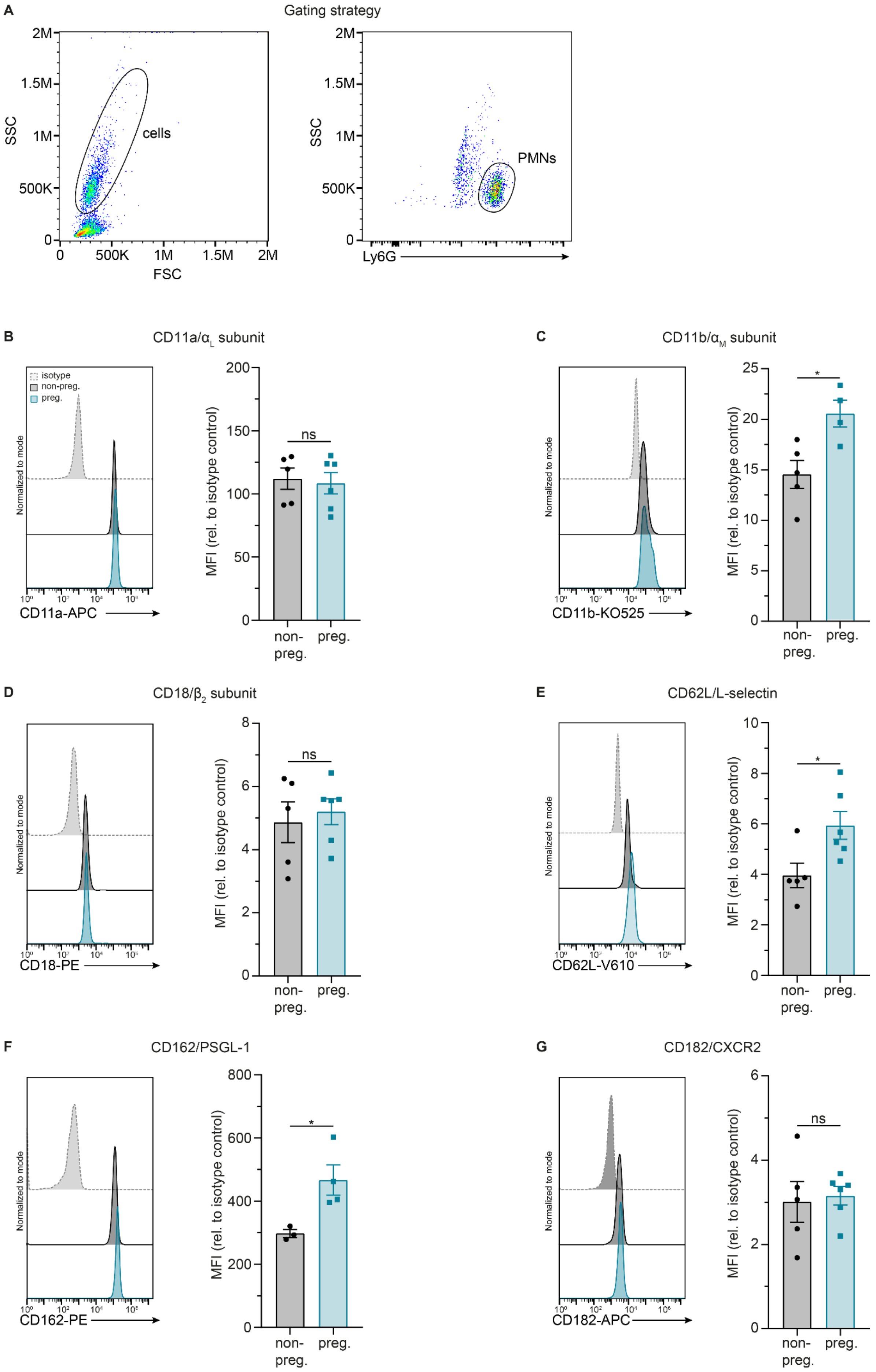

**Supplemental Figure S1 Recruitment relevant surface molecules are altered on peripheral blood neutrophils from pregnant mice.** (**A**) Gating strategy of murine whole blood samples assessing surface expression of rolling and adhesion relevant molecules on neutrophils from non-pregnant (non-preg) and pregnant (preg) mice. Surface expression levels of (**B**) CD11a, (**C**) CD11b, (**D**) CD18, (**E**) L-selectin, (**F**) PSGL-1, and (**G**) CXCR2 (n≥3mice per group; unpaired Student's t-test). Data is presented as representative flow cytometry plots and mean±SEM.

Supplemental Figure S2

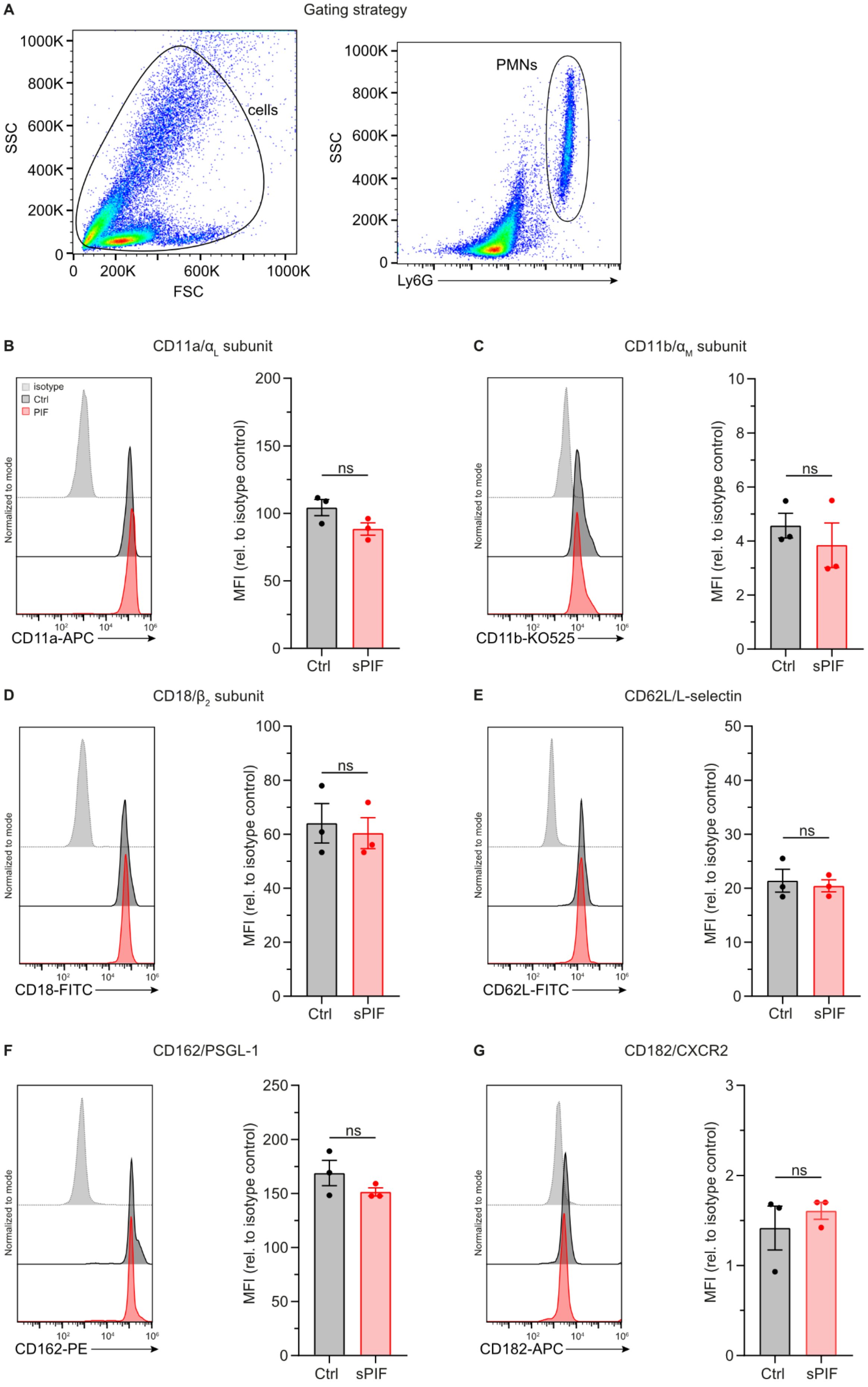

Supplemental Figure S2

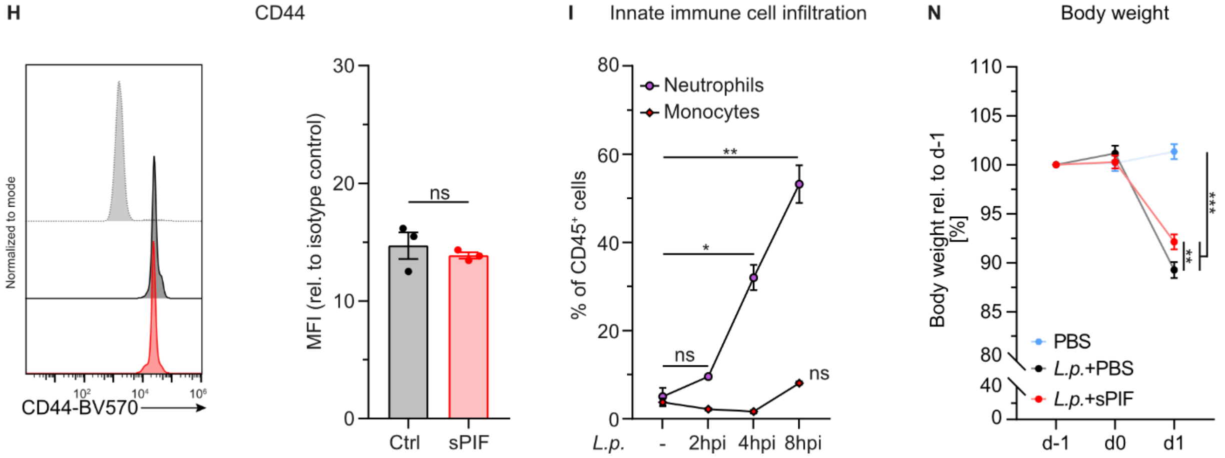


**Supplemental Figure S2 PIF does not affect surface expression of recruitment relevant surface molecules on peripheral blood neutrophils.** (**A**) Gating strategy of murine whole blood samples assessing surface expression of rolling and adhesion relevant molecules on neutrophils from WT mice injected i.s. with sPIF (1µg) or vehicle (Ctrl). Surface expression levels of (**B**) CD11a, (**C**) CD11b, (**D**) CD18, (**E**) L-selectin, (**F**) PSGL-1, (**G**) CXCR2, and (**H**) CD44. (n≥3mice per group; unpaired Student's t-test). (**I**) Infiltration of neutrophils and monocytes within 8h after *L.p.* infection (n=3 mice per group; 1-way RM ANOVA; Dunnett's multiple comparison). (**J**) Changes in body weight relative to -1dpi (n=5 mice per group; 1-way RM ANOVA; Tukey's multiple comparison). Data is presented as representative flow cytometry plots and mean±SEM.

Supplemental Figure S3

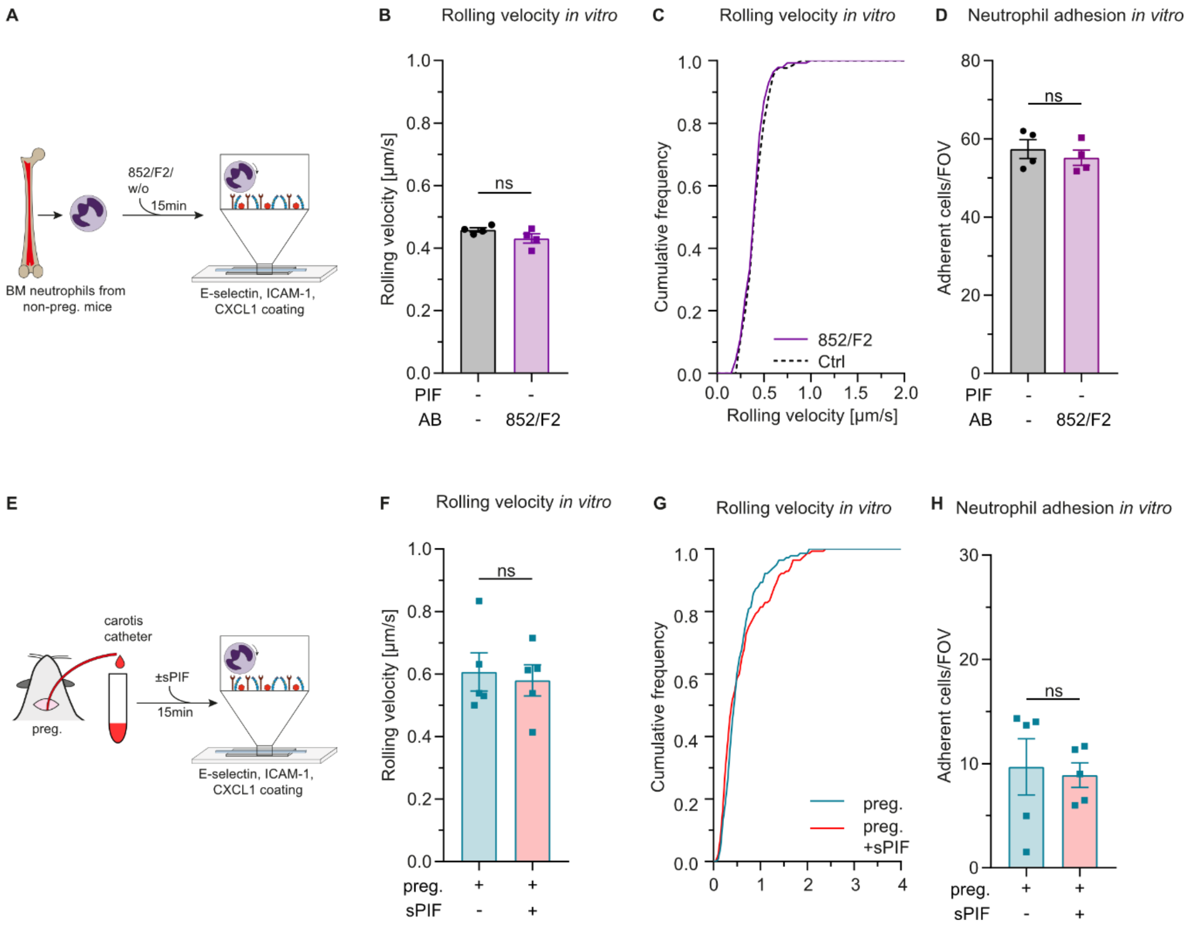


**Supplemental Figure S3 PIF-blocking antibody does not activated neutrophils and sPIF does not alter neutrophil recruitment of neutrophils from pregnant mice.** (**A**) Experimental design assessing the activating properties of PIF-blocking antibody, clone 852/F2 in E-selectin, ICAM-1 and CXCL1 coated flow chambers. (**B**) Mean rolling velocities, (**C**) cumulative distribution of rolling velocities and (**D**) number of adherent neutrophils incubated with PIF-blocking antibody (60µg $ml^{-1}$) or vehicle for 15min (n=4 independent experiments; paired Student's t-test; cumulative frequency: n=130 (w/o antibody), and n=140 (852/F2) cells). (**E**) Experimental design of flow chamber experiments assessing additional effects of recombinant PIF in whole blood from pregnant mice in E-selectin, ICAM-1 and CXCL1 coated flow chambers. (**F**) Mean rolling velocities, (**G**) cumulative distribution of rolling velocities and (**H**) number of adherent neutrophils in whole blood from pregnant mice pretreated with sPIF (300nM) or vehicle for 15min (n=5 independent experiments; paired Student's t-test; cumulative frequency: n=140 (preg), and n=140 (preg+sPIF) cells). Data is presented as mean±SEM and cumulative frequency.

Supplemental Figure S4

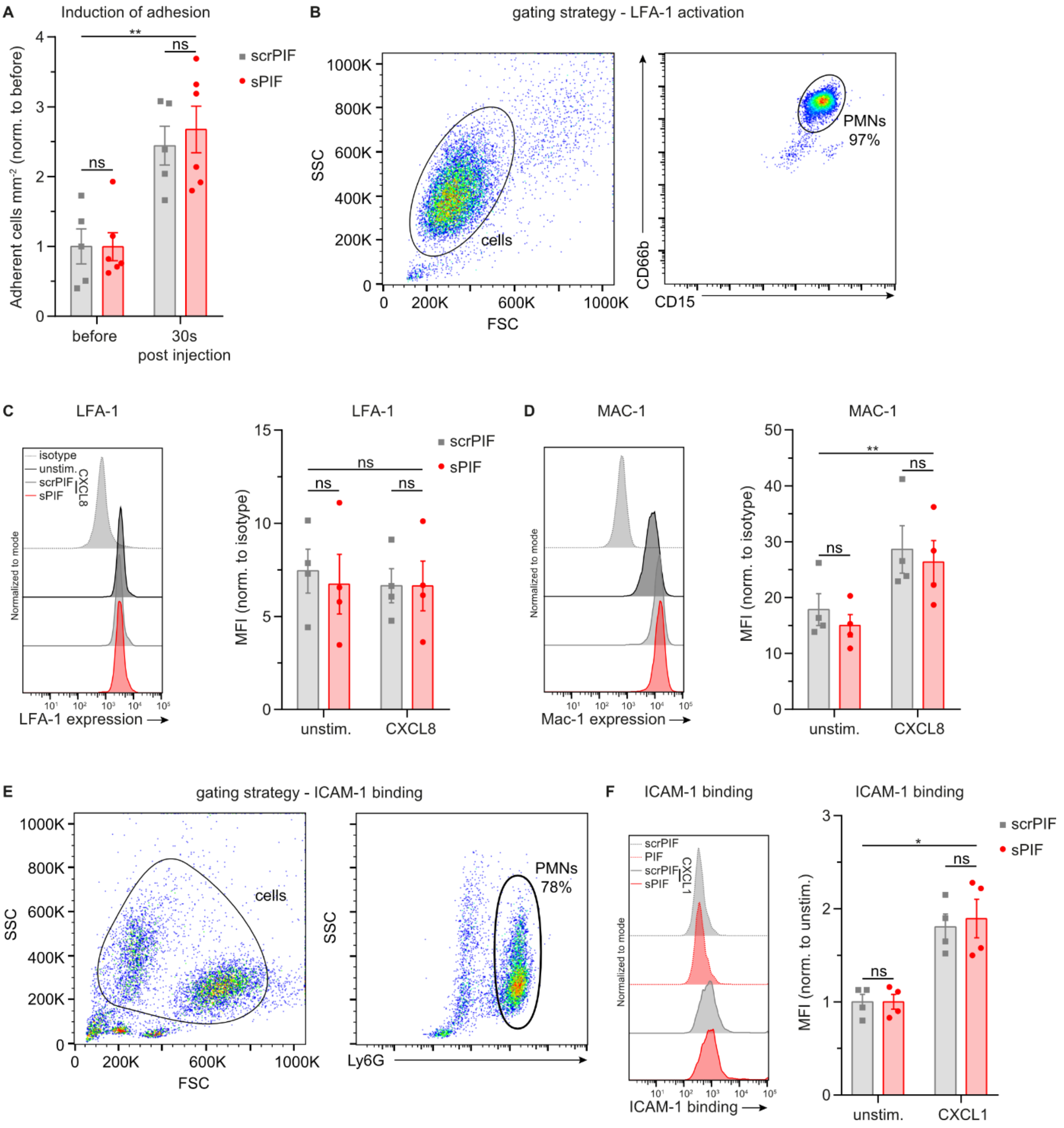


**Supplemental Figure S4 PIF does not interfere with β₂ integrin activation *in vivo* and *in vitro*.** (**A**) Induction of neutrophil adhesion (number of adherent cells mm$^{-2}$ 30s post i.a. CXCL1 injection (600ng) relative to before) in postcapillary venules of cremaster muscles of WT mice injected i.s. with sPIF or scrPIF (1µg) 2h prior to IVM (n≥5mice per group, 2-way RM ANOVA, Sidak's multiple comparison). (**B**) Gating strategy of isolated human neutrophil assessing surface expression of (**C**) LFA-1 and (**D**) MAC-1 after stimulation with CXCL8 (10nM) or vehicle (unstim.) (MFI: median fluorescence, n=4 independent experiments, 2-way RM ANOVA, Sidak's multiple comparison). (**E**) Gating strategy of isolated bone marrow neutrophils from WT mice. (**F**) ICAM-1 binding to bone marrow neutrophils pretreated with sPIF or scrPIF (300nM) upon stimulation with CXCL1 (10nM) or PBS (unstim.) for 5min (n=4mice per group; 2-way RM ANOVA, Sidak's multiple comparison). Data is presented as mean±SEM and representative flow cytometry plots.

Supplemental Figure S5

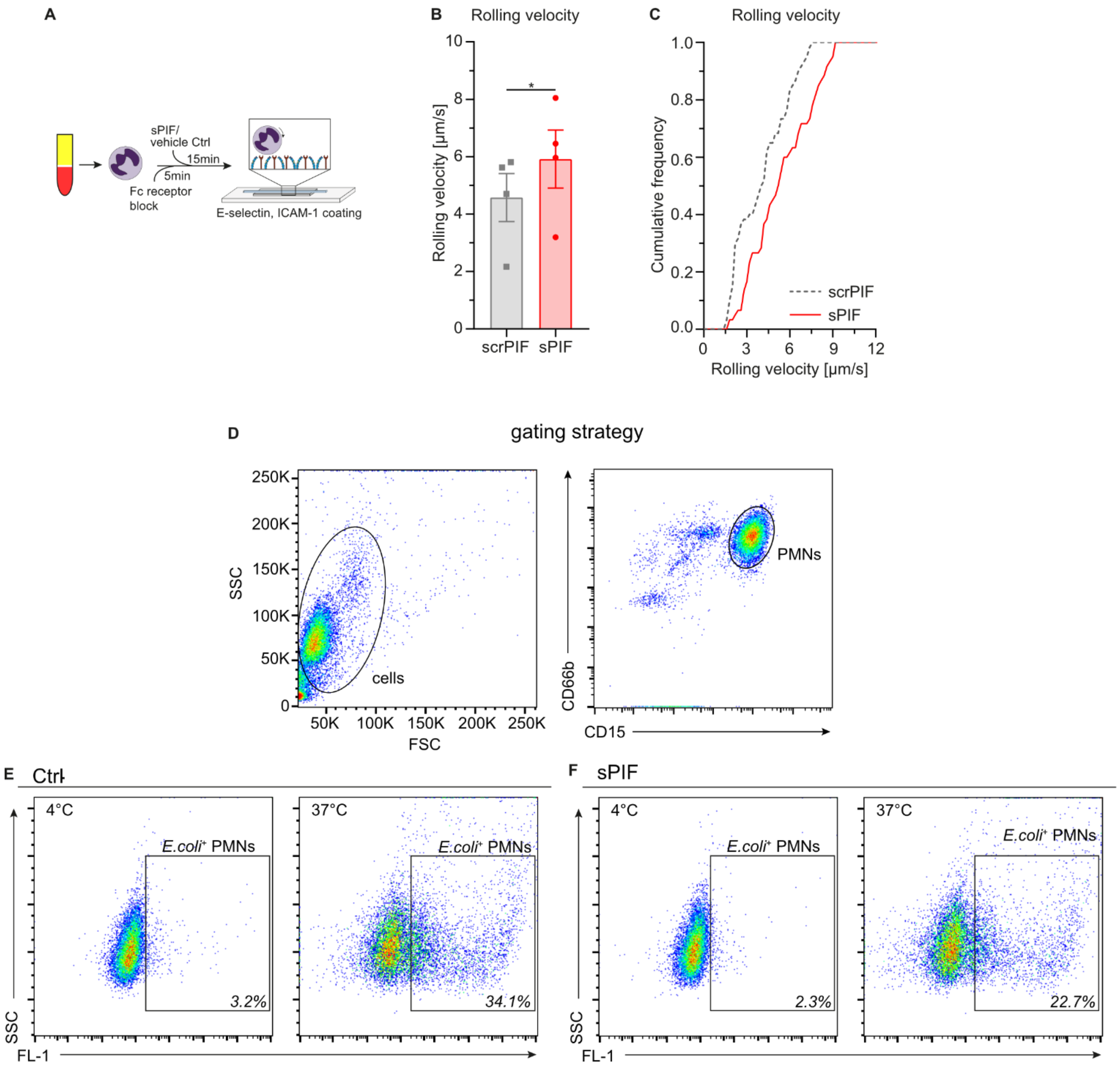


**Supplemental Figure S5 PIF impairs neutrophil adhesion *in vivo* via $K_V$1.3 and impairs neutrophil slow rolling and phagocytosis *in vitro*.** (**B**) Mean rolling velocities and (**C**) cumulative distribution of rolling velocities of isolated human neutrophils in E-selectin and ICAM-1 coated flow chambers treated with Fc block for 5min and subsequent incubation with sPIF or scrPIF (100nM) for 15min (n=4 independent experiments; paired Student's t-test; cumulative frequency: n=60 (sPIF), and n=60 (scrPIF) cells). (**D**) Gating strategy of neutrophils from human whole blood samples. Representative flow cytometry plots of (**E**) vehicle (Ctrl) and (**F**) sPIF (300nM) pretreated human whole blood incubated with fluorescent *E.coli* particles at 4°C (negative control) or 37°C. Data is presented as mean±SEM, cumulative frequency or representative flow cytometry plots.

**Supplementary Table S1 Microvascular parameters of intravital experiments of mouse mesentery.** Vessel diameter, blood flow velocity, wall shear rate, white blood cell counts (WBC), and neutrophil counts (PMN) of TNF stimulated non-pregnant (non-preg.) and pregnant (preg.) *Catchup*$^{IVM-red}$ mice (mean±SEM; unpaired Student's t-test).

| | n (mice) | n (venules) | Diameter [µm] | Centerline velocity [µm $s^{-1}$] | Wall shear rate [$s^{-1}$] | WBC [µ$l^{-1}$] | PMN [µ$l^{-1}$] |
|---|---|---|---|---|---|---|---|
| **non-preg.** | 6 | 31 | 133±5 | 3616±210 | 689±45 | 1172±107 | 297±65 |
| **preg.** | 6 | 27 | 135±5 | 3992±233 | 753±53 | 872±175 | 172±46 |
| | | | ns. (p=0.7863) | ns. (p=0.2340) | ns. (p=0.3581) | ns. (p=0.1731) | ns. (p=0.2149) |

**Supplementary Table S2 Microvascular parameters of TNF stimulated mice.** Vessel diameter, centerline velocity, wall shear rate, white blood cell counts (WBC), and neutrophil (PMN) counts of TNF-α stimulated *WT* mice pretreated with either vehicle (Ctrl), scrPIF or sPIF, respectively (mean±SEM; 1-way ANOVA, Tukey's multiple comparison).

| | n (mice) | n (venules) | Diameter [µm] | Centerline velocity [µm $s^{-1}$] | Wall shear rate [$s^{-1}$] | WBC [µ$l^{-1}$] | PMN [µ$l^{-1}$] |
|---|---|---|---|---|---|---|---|
| **Ctrl** | 6 | 14 | 33±1 | 1714±179 | 1315±151 | 3754±597 | 1513±281 |
| **scrPIF** | 6 | 21 | 33±1 | 1671±164 | 1243±117 | 4422±560 | 1645±461 |
| **sPIF** | 6 | 18 | 32±1 | 1667±196 | 1276±141 | 4605±358 | 1793±222 |
| | | | ns. (p=0.8735) | ns. (p=0.9819) | ns. (p=0.9349) | ns. (p=0.4979) | ns. (p=0.8523) |

**Supplementary Table S3 Microvascular parameters before and after CXCL1 injection.** Vessel diameter, centerline velocity, wall shear rate, white blood cell counts (WBC), and neutrophil (PMN) counts of unstimulated *WT* mice pretreated with either scrPIF or sPIF, respectively before and after injection of CXCL1 (mean±SEM; unpaired Student's t-test).

| Time | | n (mice) | Diameter [µm] | Centerline velocity [µm $s^{-1}$] | Wall shear rate [$s^{-1}$] | WBC [µ$l^{-1}$] | PMN [µ$l^{-1}$] |
|---|---|---|---|---|---|---|---|
| **before** | **scrPIF** | 5 | 32±1 | 1940±336 | 1505±256 | 6428±673 | 2588±376 |
| | **sPIF** | 6 | 30±1 | 2133±305 | 1768±289 | 7173±623 | 3018±846 |
| | | | ns. (p=0.4086) | ns. (p=0.6798) | ns. (p=0.5213) | ns. (p=0.4377) | ns. (p=0.6755) |
| **0.5min/1min** | **scrPIF** | | | 2060±441 | 1611±346 | | |
| | **sPIF** | | | 2300±388 | 1937±371 | | |
| | | | | ns. (p=0.6914) | ns. (p=0.5430) | | |
| **5min** | **scrPIF** | | | 1580±396 | 1230±301 | 3938±368 | 1734±571 |
| | **sPIF** | | | 2400±383 | 2020±370 | 4487±470 | 2132±716 |
| | | | | ns. (p=0.1728) | ns. (p=0.1420) | ns. (p=0.3970) | ns. (p=0.6834) |

**Supplementary Table S4 Microvascular parameters.** Vessel diameter, centerline velocity, wall shear rate, white blood cell counts (WBC), and neutrophil (PMN) counts of TNF-α stimulated *WT* or *Kcna3*$^{-/-}$ mice pretreated with sPIF, PAP-1, or sPIF and PAP-1, respectively (mean±SEM; 1-way ANOVA, Tukey's multiple comparison).

| | n (mice) | n (venules) | Diameter [µm] | Centerline velocity [µm $s^{-1}$] | Wall shear rate [$s^{-1}$] | WBC [µl$^{-1}$] | PMN [µl$^{-1}$] |
|---|---|---|---|---|---|---|---|
| ***WT*** | 4 | 13 | 33±1 | 1708±202 | 1265±135 | 3853±543 | 1708±520 |
| ***WT+sPIF*** | 6 | 19 | 32±1 | 1616±146 | 1268±119 | 4018±282 | 2355±350 |
| ***WT*+PAP-1+sPIF** | 4 | 13 | 29±1 | 1577±175 | 1322±131 | 3393±495 | 1670±425 |
| **WT+PAP-1+scrPIF** | 4 | 13 | 29±1 | 1669±220 | 1376±161 | 3060±322 | 1413±504 |
| ***Kcna3*$^{-/-}$+sPIF** | 5 | 19 | 31±1 | 1900±106 | 1541±106 | 2956±348 | 1386±161 |
| | | | ns. (p=0.1578) | ns. (p=0.6176) | ns. (p=0.4613) | ns. (p=0.2221) | ns. (p=0.3483) |
